# Understanding early HIV-1 rebound dynamics following antiretroviral therapy interruption: The importance of effector cell expansion

**DOI:** 10.1101/2024.05.03.592318

**Authors:** Tin Phan, Jessica M. Conway, Nicole Pagane, Jasmine Kreig, Narmada Sambaturu, Sarafa Iyaniwura, Jonathan Z. Li, Ruy M. Ribeiro, Ruian Ke, Alan S. Perelson

## Abstract

Most people living with HIV-1 experience rapid viral rebound once antiretroviral therapy is interrupted; however, a small fraction remain in viral remission for an extended duration. Understanding the factors that determine whether viral rebound is likely after treatment interruption can enable the development of optimal treatment regimens and therapeutic interventions to potentially achieve a functional cure for HIV-1. We built upon the theoretical framework proposed by Conway and Perelson to construct dynamic models of virus-immune interactions to study factors that influence viral rebound dynamics. We evaluated these models using viral load data from 24 individuals following antiretroviral therapy interruption. The best-performing model accurately captures the heterogeneity of viral dynamics and highlights the importance of the effector cell expansion rate. Our results show that post-treatment controllers and non-controllers can be distinguished based on the effector cell expansion rate in our models. Furthermore, these results demonstrate the potential of using dynamic models incorporating an effector cell response to understand early viral rebound dynamics post-antiretroviral therapy interruption.

## Introduction

Antiretroviral therapy (ART) is successful at suppressing human immunodeficiency virus-1 (HIV-1) below the limit of detection in people with HIV-1 (PWH). However, ART does not completely eliminate HIV-1 due to the presence of latently infected cells, which have low or no HIV-1 gene expression, making it difficult for the immune system to recognize and eliminate them [1]. While continual use of ART robustly controls HIV-1, there are a number of issues with long-term ART, ranging from how well PWH adhere to treatment [2–5] to the potential side effects of taking ART long term [6], such as chronic inflammation and HIV-associated neurocognitive disorders [7–10]. Thus, durable ART-free virological control remains a major goal.

Due to the persistence of the latent reservoir [11–18], if ART is interrupted, latently infected cells that reactivate can lead to viral rebound. In fact, following analytical treatment interruption (ATI) of ART, most persons with HIV-1, including those with an extremely small latent reservoir, experience viral rebound typically within weeks [19–23]. But in rare cases, some individuals maintained low viral loads for an extended duration of months to years [16,22,24,25]. Individuals, who experience rapid viral rebound, are generally referred to as non-controllers (NC), while those who control are termed post-treatment controllers (PTC). The generation of post-treatment control appears to be affected by external factors such as the timing of ART initiation [17,22,25–28], and in human and nonhuman primate models by viral suppression mediated by CD8+ cells [29–34]. Thus, understanding the factors associated with PTC can suggest a clinical path towards durable control in PWH.

Several models have been proposed to explain the mechanisms behind viral rebound following ATI. The studies by Hill et al. [35,36], Pinkevych et al. [37,38], Fennessey et al. [39], and van Dorp et al. [40] hypothesized rebound as a stochastic process due to the reactivation of latently infected cells that release virus and initiate a chain of other successful infection events. Yet, in PTC, viral load is kept at a low level despite a large reservoir size in some people. This suggests that other factors, such as the immune response, also play important roles in determining rebound dynamics of HIV-1 [41–48]. For this reason, Conway and Perelson [49] proposed a model of viral rebound considering both the latent reservoir and immune response dynamics. Their model demonstrated the combined impact of the immune response and the size of the latent reservoir on HIV-1 dynamics post-ATI and lays the foundation for this study.

Some modeling studies have also estimated the distribution of time to rebound. For instance, Conway et al. [50,51] utilized personal biomarkers including HIV-1 reservoir quantification and cell associated HIV-1 RNA to look at the distribution of time to rebound and fit a model to viral rebound data from a collection of ACTG (Advancing Clinical Therapeutics Globally for HIV/AIDS, formerly known as the AIDS Clinical Trials Group) ATI studies [16]. Others studied mathematical and statistical models to fit rebound data of HIV-1 and simian immunodeficiency virus [50,52–54], including studies in which additional treatments with monoclonal antibodies or immune stimulants were given prior to ATI [31,55,56]. While these attempts provide valuable insights into the rebound dynamics post-ATI, they either did not aim to establish the biological factors that distinguish PTC and NC or did not fit to individual viral rebound data. In this work, our objective is to examine whether mechanistic models with virus-immune interactions can accurately capture the heterogeneity in viral rebound dynamics and identify potential mechanisms that distinguish the outcomes post-ATI.

## Method

### Ethics Statement

This research was approved by the LANL Human Subjects Research Review Board (HSRRB).

### Data

We analyze the data of 24 PWH (9 PTC and 15 NC) from Sharaf et al. [57] for whom we have longitudinal viral load data. The PTC participants were identified from several clinical studies in the ACTG [58–62]. In Sharaf et al., PTCs were defined as individuals who maintained a viral load of less than 400 HIV-1 RNA copies/mL for at least 24 weeks post ATI and where short-term increases (≤ 2 viral load measurements) of over 400 HIV-1 RNA copies/mL were not exclusionary. Non-controllers were defined as individuals who did not meet the PTC definition. We chose to analyze the data between 0- and 52-weeks post ATI to focus on the period during which the viral load data is often used to classify PTC and NC in clinical settings.

### Mathematical models

Conway and Perelson [49] introduced the following model of viral dynamics to study the phenomenon of PTC:

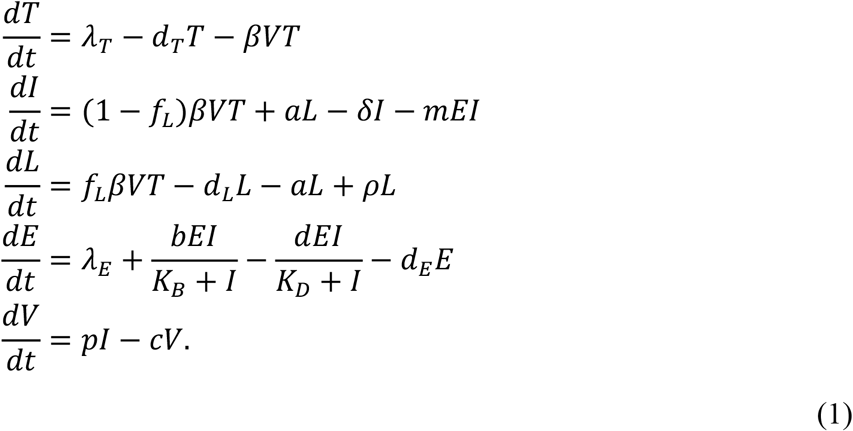

This model (Eqns. 1) assumes that target cells (*T*) are produced and die at constant rates *λ*_*T*_ and *d*_*T*_, respectively. Virus (V) infects target cells with rate constant *β* and is cleared at per capita rate *c*. The infection leads to productively infected cells (*I*) at a rate *f*_*I*_*βVT*, where *f*_*L*_ and (1 ™ *f*_*L*_) are the fractions of infections that lead to latently and productively infected cells, respectively. Infected cells release viruses at per capita rate *p* and die at per capita rate *δ* by viral cytopathic effects. Latently infected cells (*L*) proliferate and decay at per capita rates *ρ* and *d*_*L*_, respectively, and can reactivate at per capita rate *a* to become productively infected cells. In the absence of viral infection, effector cells (*E*), such as CD8+ T-cells and natural killer (NK) cells, are produced at rate *λ*_*E*_ and die at per capita rate *d*_*E*_. During viral infection, the effector cells recognize infected cells, which stimulates its population to expand. Conway and Perelson modeled this process using the infected-cell dependent growth term 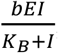, where *b* is the maximal growth rate and *K*_*B*_ is the density of infected cells required for the effector cell growth rate to be at its half-maximal rate. Note if *K*_*B*_ is very large, then a large number of infected cells will be needed to activate the expansion of effector cells, and this will lead to a less effective effector cell response.

The effector cells kill productively infected cells with rate constant *m* and can become exhausted at maximal rate *d*. Effector exhaustion is induced by the upregulation of inhibitory receptors during effector expansion that ultimately abrogate the expansion and other functional responses. Chronic infections and cancers typically lead to effector cell exhaustion, which occurs through metabolic and epigenetic reprogramming of the effector cells that incrementally make them worse at proliferating, surviving, and performing effector functions, such as killing and releasing diffusible molecules [63–65]. Conway and Perelson modeled the rate of exhaustion with a similar infected-cell dependent functional response term 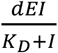. These functional response terms had previously been suggested by Bonhoeffer et al. [66].

The model formulation assumes a fraction, *f*_*L*_, of infections lead to the production of latently infected cells. The homeostatic proliferation of latently infected cells at per capita rate ρ follows from prior work [67,68] and is supported by experimental evidence showing that latent cells can proliferate without activating (or expressing viral signals) [18,69–72].

*A Simplified Model*. We formulated variations of the Conway and Perelson model by altering the model assumptions on the dynamics of latent cells and the effector cells (S1, Supplementary Material). In particular, without effector cell exhaustion, we have Simplified Model 1:

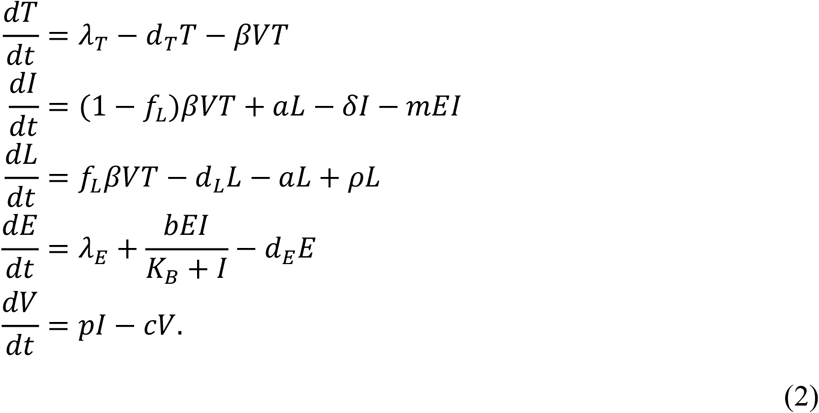

For further simplification, we assume that the number of viruses is in quasi-steady state with the number of infected cells in all models including the Conway and Perelson model, so that 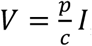 as in prior work [73,74]. The initial time (*t* = 0) is when the individuals are taken off ART. We 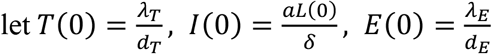, and 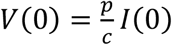, which are approximations of the steady states during ART assuming a small latent reservoir at the time of ATI, that ART was highly effective and suppressed all new infections, and thus that productively infected cells were only generated via activation of latently infected cells. We also studied other models given in the Supplementary Material. In models without explicit latently infected cell dynamics, we assume the reservoir size does not vary significantly during the first year post-ATI or during the off-ART duration for NCs, the time period we study. This implies *L*(*t*) = *L*(0). Whether the viral load rebounds or not post-ATI is determined by both the immune response and the latent reservoir size. The influence of the reservoir size at the time of ATI has been studied before [36,49], thus we choose to examine whether the immune response — without variation in the latent reservoir size — can capture the viral rebound dynamics for this cohort of participants.

The parameters that we fit, and the fixed parameters are given in Table 1. *f*_*L*_ is fixed to 10^−6^, meaning only 0.0001% of infections result in latent cell production [75]; however, increasing *fL* up to 10-fold, has little effect on the model dynamics. The assumed size of the latent reservoir was chosen as 1 cell/ml based on estimates of the number of replication competent latently infected cells per million resting CD4 T cells determined by the quantitative viral outgrowth assay [76,77] and assuming patients on long-term ART have close to a million CD4+ T cells per ml. The net rate of change of the latent cell population *d*_*L*_ − *a* + *ρ* is such that the expected half-life (*t*_1/2_) of the reservoir is 44 months [14,15]. We set *d*_*L*_ = 0.004 per day [68], and *a* = 0.001 [49], which gives *ρ* ≈ 0.0045 per day. We remark that the stability of the latent reservoir is due to the combined effect of proliferation, death, and activation; however, these parameter values are only educated guesses. Furthermore, recent data has suggested that due to proliferation, the reservoir might stop decaying [18].

**Table 1.** Model parameters. The double dashed line separates parameters that we fit (above) and fixed (below). References for the fitting parameters are provided for comparison purposes.

| *Parameter* | *Meaning* | *Unit* |  |
| --- | --- | --- | --- |
| $\beta$ | Infection rate | $mL (HIV RNA copies)^{-1} day^{-1}$ | [78,79] |
| $\lambda_E$ | Production rate of effector cells | $cells mL^{-1} day^{-1}$ | [49] |
| $m$ | Effector cell killing rate | $mL cells^{-1} day^{-1}$ | [49] |
| $b$ | Maximum effector cell growth rate | $day^{-1}$ | [80] |
| $K_B$ | Density of infected cells needed to reach a half-maximal effector cell growth rate | $cells mL^{-1}$ | [66] |
| $p$ | Viral production rate | $(HIV RNA copies) day^{-1}$ | [49] |
| $d$ | Maximum effector cell exhaustion rate | $day^{-1}$ | [81] |
| $K_D$ | Density of infected cells needed to reach a half-maximal effector cell exhaustion rate | $cells mL^{-1}$ | [66] |
| $\lambda_T$ | Production rate of target cells | $10^4 cells mL^{-1} day^{-1}$ | [82] |
| $d_T$ | Death rate of target cells | $0.01 day^{-1}$ | [68] |
| $f_L$ | Fraction of latently infected cells produced per infection | $10^{-6}$ | [49] |
| $a$ | Reactivation rate of latently infected cells | $0.001 \text{ day}^{-1}$ | [49] |
| $\delta$ | Death rate of infected cells | $1 \text{ day}^{-1}$ | [83] |
| $d_E$ | Death rate of effector cells | $2 \text{ day}^{-1}$ | [81] |
| $L_0$ | Steady state size of the latent reservoir | $1 \text{ cell mL}^{-1}$ | [76,77] |
| $d_L$ | Decay rate of the latent reservoir | $0.004 \text{ day}^{-1}$ | [14,68] |
| $\rho$ | Latently infected cell proliferation rate, $d_L + a - \log(2)/t_{1/2}$ | $0.0045 \text{ day}^{-1}$ | [49,84] |
| $c$ | Clearance rate of virus | $23 \text{ day}^{-1}$ | [85] |

### Data Fitting

We used a nonlinear mixed effects modeling approach (software Monolix 2023R1, Lixoft, SA, Antony, France) to fit Simplified Model 1 and other variants presented in the Supplementary Material to the viral load data for all individuals simultaneously. We applied left censoring to data points under the limit of detection.

We assumed the fitting parameters follow a logit-normal distribution as we constrained the parameters to be in a fixed range. *λ*_*E*_, *b, m*, and *d* were constrained between 0 and 10 times the reference values used in Conway and Perelson (Table 1). Imposing bounds on *λE, b, m*, and *d* is not necessary; however, this allows us to get the most consistent results across all models tested.

We fit *β* and constrained its range between 10^-7.5^ and 10^−12^ mL (HIV RNA copies)^-1^ day^-1^ to avoid an unrealistically high viral surge within 24 hours post-ATI due to high values of *β*. Parameter *p* was constrained between 0 and 5000 per day. Parameters *K*_*B*_ *and K*_*D*_ were constrained between 0 and 100,000 cells per mL. Without this upper bound, *K*_*D*_ value often tended to infinity. No covariate was used during the initial fitting and comparison of all models. A covariate based on whether a participant is classified as PTC or NC was used later with the best fit model to determine the factors that can distinguish these two groups. Model comparison was done using the corrected Bayesian Information Criterion (BICc) [86] as reported by Monolix.

## Result

### Model fit and comparison

We fit the Conway and Perelson model and variations of it with different assumptions on the dynamics of latent and effector cells (Supplemental Material). The best fit model was Simplified Model 1 described in Eq. (2), where effector cell exhaustion is disregarded. Its best-fit to the data is shown in Figure 1. Comparisons of best fits for other models along with population and individual parameters are shown in Figures S1-S6 and Tables S1-S13. The only difference between Simplified Model 1 and the Conway & Perelson model is the lack of an exhaustion term, which suggests exhaustion is not necessary to explain early viral rebound dynamics (within the first 52 weeks) seen in this set of PWH. This is further supported by the negligible effect of exhaustion estimated from the best fits for the Conway & Perelson model and Simplified Model 3, where the values for *d* and *K*_*D*_ result in a much smaller exhaustion effect compared to the reference values (Tables S2, 4). This does not imply that effector exhaustion is not relevant to HIV rebound dynamics as it is potentially important to explain the eventual loss of control as occurred in the subset of PTCs who have a truncated x-axis in Figure 1 and in the CHAMP study [25]. In addition, it is an important mechanism to explain the phenomenon where a PTC can have a high viral set point pre-ART and a lower viral set point post-ATI [49], and a variety of other dynamical phenomena [56,64,87].

**Figure 1.**
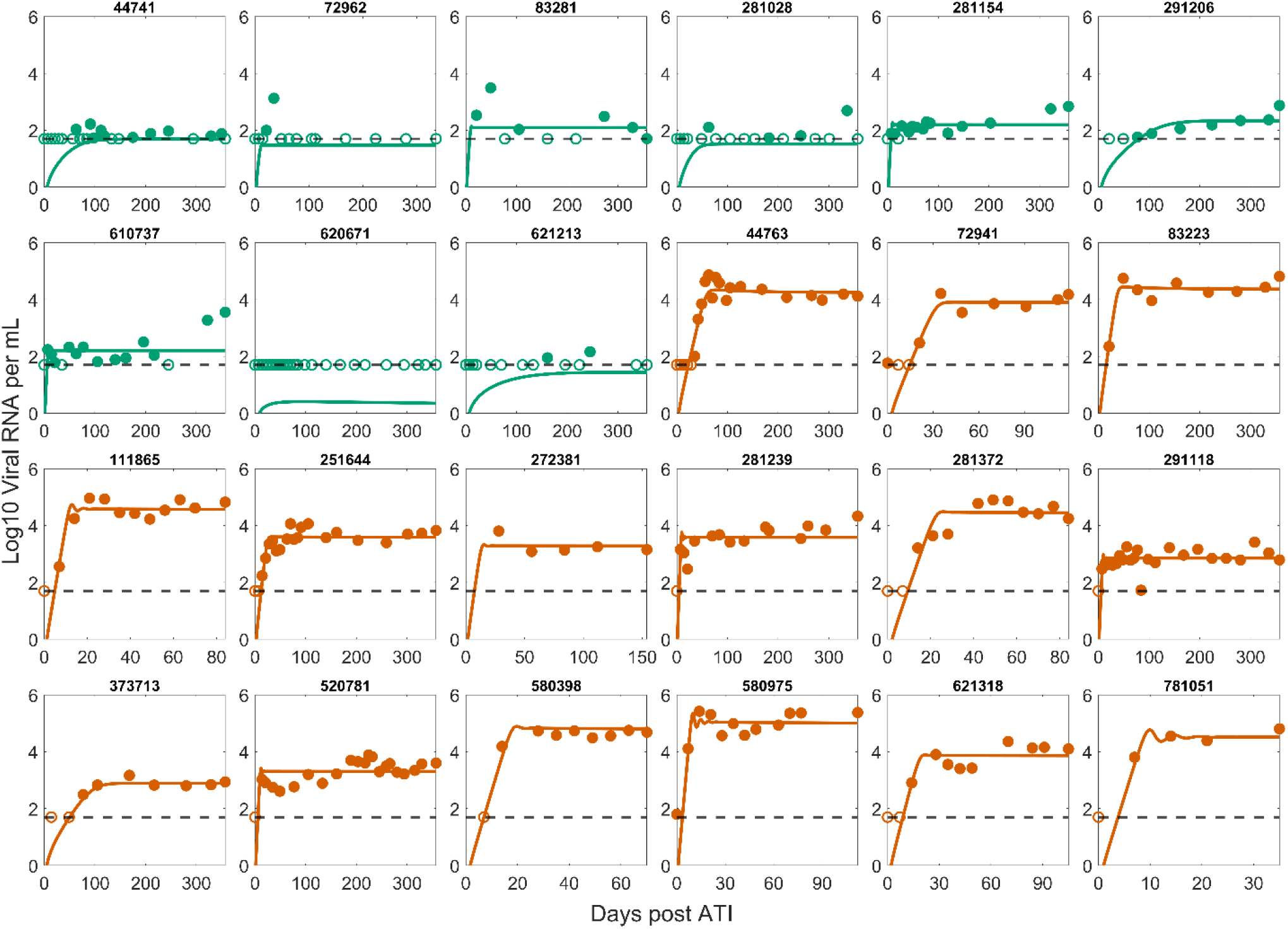
Best fit of Simplified Model 1 to the post-ATI data from Sharaf et al. [57]. Green indicates PTC. Dark orange indicates NC. The horizontal dashed line is the limit of detection. Open circles are data points below the limit of detection (50 viral RNA copies/mL). Filled circles are data points above the limit of detection. Note the x-axis scale varies for the NCs as many of them were put back on ART terminating the data set.

Interestingly, the best fit parameter values for Simplified Model 1 with the exception of *K*B are close to the parameter values used in Conway and Perelson (compare Tables 1 and 2). This is also true for Simplified Model 1 even when we do not impose bounds on *λ*_*E*_, *b*, and *m* (not shown). All models with effector cell dynamics fit the data relatively well (Table S1), which is indicative of the importance of including in the model the dynamics of a population of effector cells that can kill productively infected cells.

**Table 2.** Best fit population parameters (Simplified Model 1) vs. reference values from Conway and Perelson [49].

| Parameter | Best fit values (population estimate) | Reference values |
| --- | --- | --- |
| $\beta$ | $1.51 \times 10^{-8} \text{ mL day}^{-1}$ | $1.5 \times 10^{-8} \text{ mL day}^{-1}$ |
| $\lambda_E$ | $1.45 \text{ mL}^{-1} \text{ day}^{-1}$ | $1 \text{ mL}^{-1} \text{ day}^{-1}$ |
| $m$ | $0.83 \text{ mL day}^{-1}$ | $0.42 \text{ mL day}^{-1}$ |
| $b$ | $4.68 \text{ day}^{-1}$ | $1 \text{ day}^{-1}$ |
| $K_B$ | $52.62 \text{ mL}^{-1}$ | $0.1 \text{ mL}^{-1}$ |
| $p$ | $3477 \text{ day}^{-1}$ | $2000 \text{ day}^{-1}$ |

We stratified the best fit parameters (Simplified Model 1) for each person by whether they were PTC or NC (Figure S8). The most statistically significant difference using the Mann-Whitney test between the PTC and NC group is the parameter *K*_*B*_ (*p*-value: 0.0005), followed by the immune cell killing rate *m* (*p*-value: 0.0169). All model parameters differ at least somewhat between these two groups; however, the degree of significance of the difference for each parameter varies among models. In contrast, the statistically significant difference of *K*_*B*_ between PTC and NC is consistently observed across all models tested (Figures S7-S10).

This observation prompted us to fit Simplified Model 1 using a covariate on *K*_*B*_ between PTC and NC. This modification significantly improves the model by 26 BICc points (Figure S2, Table S1); however, further inclusion of other covariates improves the fit slightly but worsens the BICc score (Table S1). The best fit population value of *K*_*B*_ for PTC 1.19 cell mL^-1^ is consistent with the value in Conway and Perelson [49], which indicates a fast expansion of effector cells. The best fit population value of *K*_*B*_ for NC is 230-fold higher, indicating a slower expansion of effector cells (Table 3).

**Table 3.** Best fit population parameters (Simplified Model 1 with a covariate on K_B_) vs. reference values from Conway and Perelson [49].

| *Parameter* | *Best fit values (population estimate)* | *Reference values* |
| --- | --- | --- |
| $\beta$ | $1.58 \times 10^{-8}$<br>$\text{mL (HIV RNA copies)}^{-1}\text{day}^{-1}$ | $1.5 \times 10^{-8}$<br>$\text{mL (HIV RNA copies)}^{-1}\text{day}^{-1}$ |
| $\lambda_E$ | $0.50 \text{ cells mL}^{-1}\text{day}^{-1}$ | $1 \text{ cells mL}^{-1}\text{day}^{-1}$ |
| $m$ | $1.72 \text{ mL cell}^{-1}\text{day}^{-1}$ | $0.42 \text{ mL cell}^{-1}\text{day}^{-1}$ |
| $b$ | $5.40 \text{ day}^{-1}$ | $1 \text{ day}^{-1}$ |
| $K_B$ (PTC) | $1.19 \text{ cells mL}^{-1}$ | $0.1 \text{ cells mL}^{-1}$ |
| $K_B$ (NC) | $277 \text{ cells mL}^{-1}$ | — |
| $p$ | $3220 \text{ (HIV RNA copies) day}^{-1}$ | $2000 \text{ (HIV RNA copies) day}^{-1}$ |

Stratification of the best fit parameter values for Simplified Model 1 with a covariate on *K*_*B*_ shows that the inclusion of a covariate on *K*_*B*_ is sufficient to explain the variation between PTC and NC (Figure 2). Indeed, comparing the stratified individual parameters estimated using Simplified Model 1 without a covariate (Figure S8) and with a covariate on *K*_*B*_ (Figure 2), we find that the statistically significant difference between the PTC and NC observed for all other model parameters vanishes. For completeness, we also tested whether having a covariate on other parameters can improve the fit and remove the statistically significant variation in all other parameters including *K*_*B*_. We find that adding a covariate on other parameters (one by one) not only does not improve the fitting compared to the fit with a covariate on *K*_*B*_ (Table S1), but it also has no effect on the statistically significant difference in *K*_*B*_ between PTC and NC (not shown). To see whether the model can fit the data well without accounting for individual variation in *K*_*B*_, we tested Simplified Model 1 without random effect for *K*_*B*_, which results in worse fits (Table S1). Altogether, these results support the inclusion of a covariate on *K*_*B*_ as well as including individual variation in *K*_*B*_.

**Figure 2.**
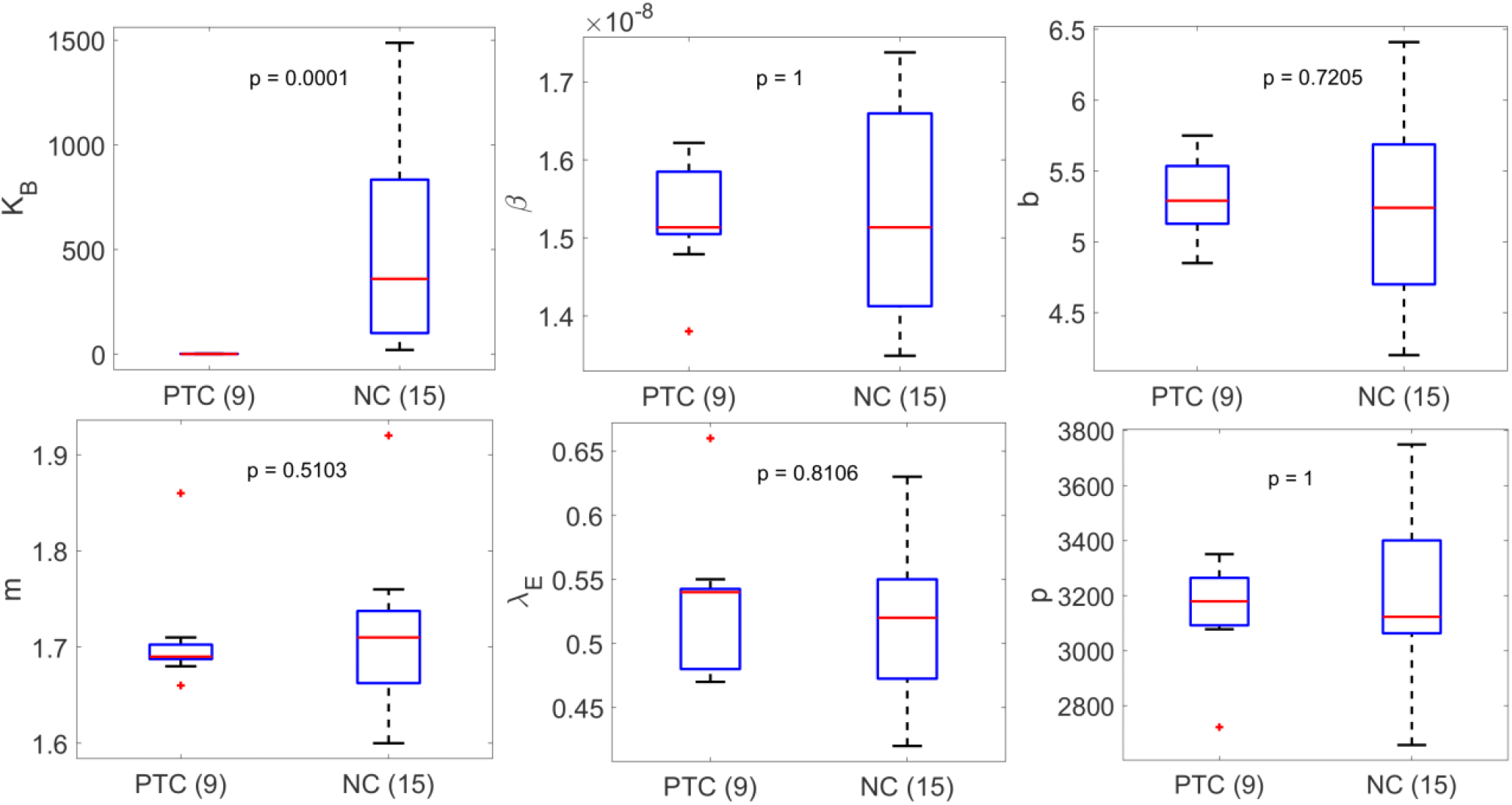
Summary of best fit parameters in Simplified Model 1 with a covariate on K_B_ stratified based on PTC or NC. The median value of K_B_ for PTC is 0.94 cell mL^-1^. The difference in the values of K_B_ is the most significant between PTC and NC. All other parameters are not statistically different between PTC and NC.

### Analytical approximation of the rebound dynamics

To better explain why *K*_*B*_ separates the NC and PTC participants, we approximated analytically the viral set point *V*_*ss*_ after rebound (see Supplementary Material) and found

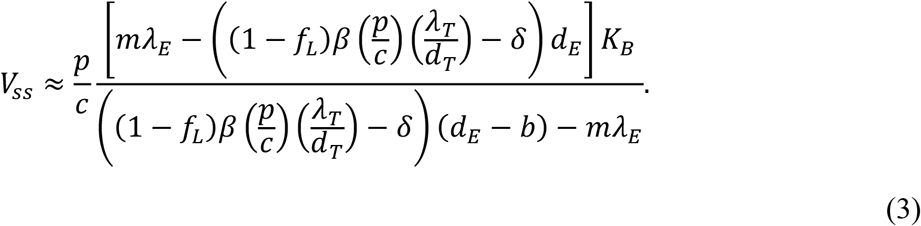

Comparing *V*_*ss*_ with the predicted viral load set-point obtained with the full model dynamics (Figure S11) demonstrates the accuracy of this approximation.

The clinical classification of PTC individuals by Sharaf et al. is centered on the viral set point and how long it takes to rebound above the threshold of 400 viral RNA copies/mL (i.e., 24 weeks or longer of viral load below 400 copies/mL). The viral set point approximation *V*_*ss*_ shows that the rebound classification is influenced by many factors in highly nonlinear ways. Yet, *V*_*ss*_ is simply proportional to *K*_*B*_, which may explain why the estimated value of *K*_*B*_ for each individual separates PTC and NC in the model fits (Tables S8).

### Distinct immune response between PTC and NC

We calculated the effector cell expansion rate per infected cell 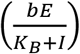 and the ratio of effector cells to infected cells (*E*/*I*) using the best fit parameters for each individual given in Table S9. There is a clear distinction in both quantities between PTC and NC (Figure 3). The effector cell expansion rate for individuals in the PTC group is several times higher than for individuals in the NC group (Figure 3A). This result is not surprising given that *K*_*B*_ is the only statistically significant difference between the two groups, where higher *K*_*B*_ for the NC group leads to a slower expansion rate of the effector cells. Additionally, in PTC individuals, the model predicts at least one effector cell per infected cell throughout the studied duration, whereas in NC individuals, the model predicts about one effector cell for hundreds of infected cells on average (Figure 3B).

**Figure 3.**
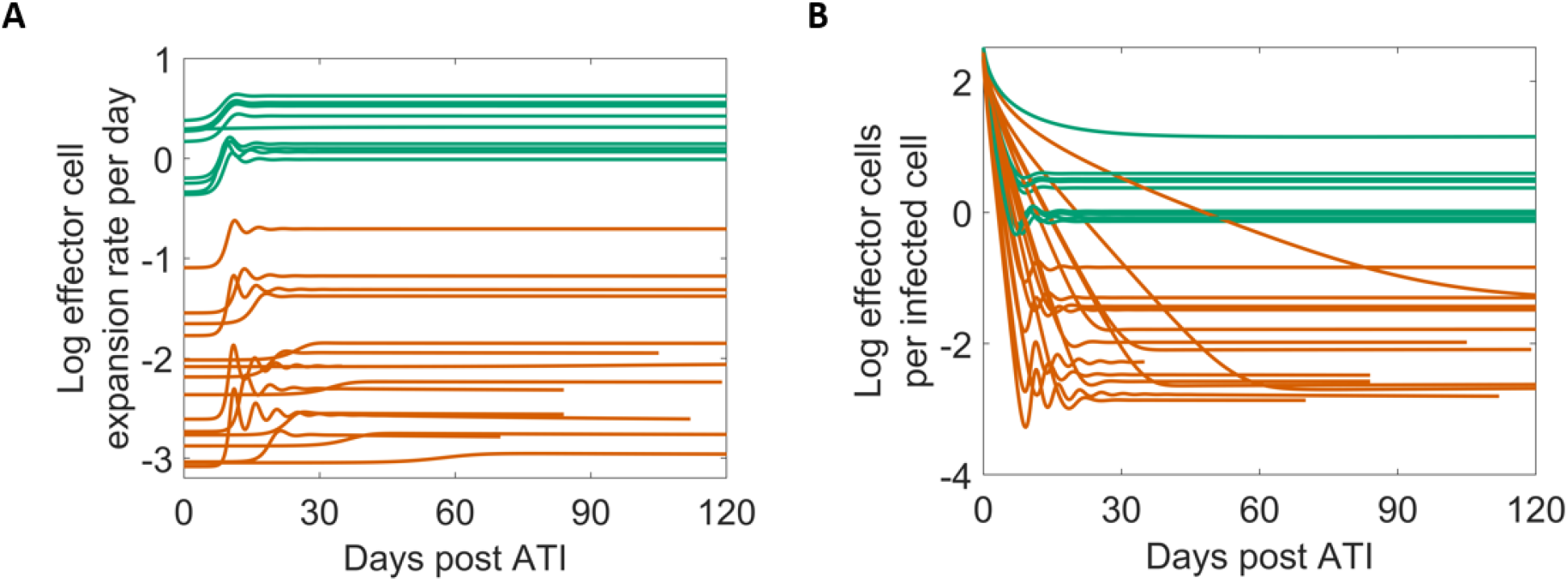
Model predicted viral and immune dynamics using Simplified Model 1 with a covariate on K_B_. Green is PTC and dark orange is NC. (A) The effector cell expansion rate 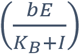 on a log10 scale. (B) The ratio of E vs. I plotted on a log10 scale.

## Discussion

Differences in the size of the latent reservoir at the time of ART interruption and virus-immune interactions have been recognized as key factors in achieving post-treatment control [49]. Here, our modeling results robustly demonstrate that differences in the expansion rate of the effector cells differentiate PTC and NC in this cohort of 24 PWH.

The size of the latent reservoir determines the overall rate of reactivation of latent cells, which can lead to HIV-1 rebound post-ATI, as pointed out by Hill et al. [35]. However, to do so requires the reactivated cells to start a chain of infections that leads to exponential growth. If the immune system detects these newly reactivated cells and reacts fast enough and strongly enough, it can interrupt this chain of infection events and lead to PTC rather than non-control. Thus, in addition to the size of the reservoir, which Sharaf et al. [57] found to be 7-fold lower in PTCs than in the NCs, other factors, such as virus-immune interactions, may also affect the viral dynamics post-ATI. Esmaeilzadeh et al. [47] showed that autologous neutralizing antibodies apply selection pressure on rebounding variants and may play a role in mediating HIV suppression after ATI. Conway and Perelson [49] suggested that when the immune response is strong, PTC can be achieved. Our modeling results illustrate this concept by accurately capturing the early viral rebound dynamics in this cohort of 24 PWH.

We found that the *K*_*B*_, the density of infected cells needed to stimulate effectors cells into growth at their half-maximal rate, is significantly lower in PTC compared to NC. Since the clinical definition of PTC depends on the post-ATI set point viral load, and we showed there is an approximately linear relationship between this viral set point and *K*_*B*_ in our model (Eqn. 3), this explains the distinct estimates of *K*_*B*_ for PTCs and NCs. These results established an analytical relationship between an early and suppressive immune response and the viral rebound dynamics, echoing the recent results in the SIV system by Vemparala et al. [88]. In practice, it is unlikely that all model parameters including *K*_*B*_ for an individual can be estimated prior to an ATI study. However, if an estimate of the value of *K*_*B*_ can be obtained, perhaps via correlation with individual’s immunological markers, then it may be possible to determine a threshold value of *K*_*B*_ that separates PTCs and NCs a priori.

A plausible biological interpretation of the variation in *K*_*B*_ comes from the studies of T cell expansion after contact with an antigen presenting cell (APC) [89,90]. Derivation of *K*_*B*_ based on the binding kinetics of a T cell to an APC results in 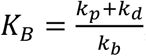, where *k*_*b*_, *k*_*d*_, and *k*_*p*_ are the rate constants for T cells binding, dissociation and first order activation/proliferation, respectively. This means that a higher binding affinity, i.e., higher *k*_*b*_ and/or lower *k*_*d*_, would result in a lower *K*_*B*_, which gives the interpretation that effector cells in PTC individuals have a higher binding affinity to the antigens presented on the surface of HIV-1 infected cells than the effector cells in NC individuals. This implies that the observed variation in *K*_*B*_ may relate to different class I HLA alleles that present the HIV-1 peptides to CD8+ T cells carried by PTC and NC [91–96]. Alternatively, *K*_*B*_ could also vary with the effector cell proliferation rate and cytotoxicity. For example, antigenic stimulation in vitro and ex vivo experiments have shown that CD8+ T cells from controllers can proliferate and increase their cytolytic capability more efficiently than those from progressors [97–99]. The effector cell population also includes other cell types, such as NK cells, and there is some evidence that NK cells may play an earlier role than CD8+ T-cells in the control of HIV-1 [42,48]. However, these interpretations remain speculations as *K*_*B*_ is modeled as the number of infected cells needed to stimulate robust effector cell expansion, so the precise mechanism associated with *K*_*B*_ is difficult to infer without a more detailed immunological model and data. Delineating the exact biological mechanisms that contribute to the differentiating values of *K*_*B*_, perhaps via ex-vivo stimulation, characterization, and TCR-sequencing of CD8+ effector cells in addition to viral sequencing from individuals participating in ATI trials, may aid in finding the right targets for an effective design of vaccines and treatments for HIV-1.

Our results demonstrate that a variation of the Conway-Perelson model without effector cell exhaustion can characterize the viral rebound dynamics in the cohort of 24 individuals from Sharaf et al. [57]. However, there are limitations to the current study that should be noted for future investigation. First, although there is considerable heterogeneity in viral rebound dynamics in the 24 studied participants, the set of possible viral rebound trajectories is more complex. For instance, the CHAMP study [25] contains individuals with more complex viral rebound dynamics and sometimes a very high initial viral peak before viral remission, which our model fits miss. In addition, since we employed a population fitting approach to fit a data set with mostly non-oscillatory individual data, the best fit model fails to capture the oscillatory features exhibited by several individuals (111865, 520781, 580975, and 621318 – Figure 1 and S2). This suggests further examination of our modeling framework using data with more heterogeneous viral rebound dynamics is necessary to generalize our results.

Second, while our parameter estimates indicate limited exhaustion of effector cells (Tables S2, 4), this could be due to several factors such as the duration of ART and the one-year time constraint we put on the data. ART has been shown to restore the functionality of HIV-1 specific CD8+ T-cells and NK cells [100,101], hence lowering the exhaustion level of effector cells. Therefore, exhaustion at the start of ATI may potentially be very limited and unobservable to our model within the one-year time constraint. This limitation is evident when consider participants such as PTC number 610737, who showed signs of viral rebound (∼2 log increase in viral load) toward the end of year one, which is not captured by the best fit models. Future studies can use longer longitudinal data to look further into the interaction of the immune system and virus over longer time scales to examine the possibility of rebound after the initial short-term viral remission. When considering a time scale on the order of years, some of the assumptions we made in order to derive the approximation of the viral set point are unlikely to hold such as that target cells remain at their disease-free equilibrium value of *λ*/*d*_*T*_ – see the derivation in S5. Thus, this necessitates more intricate mathematical analyses to ascertain the impact of immune parameters, such as *K*_*B*_, on the viral set point [102].

A third - and intriguing – observation, which we did not consider, is that in some PWH, the viral set point post-ATI is orders of magnitude lower than before ART [22]. Qualitatively, this relates to the bistability of the Conway-Perelson model with respect to different initial conditions such as the size of the latent reservoir and the immune response at the time of ATI. If ART can sufficiently reduce the viral burden, it may allow the immune response to catch up. This may happen via the restoration of functionality for HIV-1 specific CD8+ T-cells and NK cells during ART [100,101]; however, the restoration effect may not be sufficient to improve the immune response in NCs to the level of PTCs in most cases [103].

Lastly, by choosing to focus on estimating the immune response parameters in our study, we assumed that the initial conditions of the model for all 24 individuals at the time of ATI were the same. While this assumption is not ideal, when fitting both the initial conditions and the immune parameters, the resulting parameter estimates vary significantly across all participants and models, likely due to parameter identifiability issues, which makes determining the key parameters an unrealistic task. Nevertheless, even with these limitations, our modeling results robustly demonstrate that variation in the effector cell expansion rate sufficiently captures the heterogeneity in viral rebound dynamics post-ATI.

## Acknowledgement

We would like to thank Christiaan van Dorp for his insightful comments. We thank the participants, staff, and investigators of the ACTG studies.

## Disclaimer

This report was prepared as an account of work sponsored by an agency of the United States Government. Neither the United States Government nor any agency thereof, nor any of their employees, makes any warranty, express or implied, or assumes any legal liability or responsibility for the accuracy, completeness, or usefulness of any information, apparatus, product, or process disclosed, or represents that its use would not infringe privately owned rights. Reference herein to any specific commercial product, process, or service by trade name, trademark, manufacturer, or otherwise does not necessarily constitute or imply its endorsement, recommendation, or favoring by the United States Government or any agency thereof. The views and opinions of authors expressed herein do not necessarily state or reflect those of the United States Government or any agency thereof.

## Data Availability

All data are available upon request from with the written agreement of the AIDS Clinical Trials Group.

## Supplementary Material

### S1 Equations for each model

Here, we present the equations for all six models that we fit to the viral load data for the 24 PWH that we studied in the main text. Note that 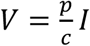 in all models.

Conway & Perelson model

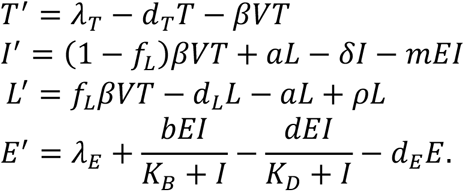

Simplified Model 1 (as above, but *d* = 0)

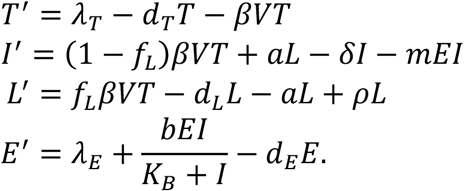

Simplified Model 2 (as Simplified Model 1, but no dynamics of latent cells *L*^*’*^ ≡ 0)

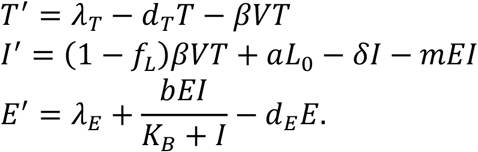

Simplified Model 3 (as Simplified Model 2, but *d* ≠ 0, so with exhaustion of effector cells)

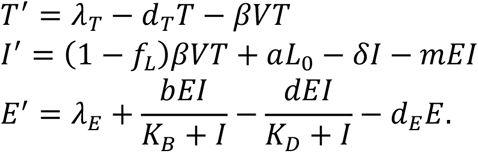

Simplified Model 4 (as Simplified Model 2, but no explicit dynamics of effector cells *E*^*’*^ ≡ 0)

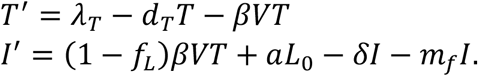

Simplified Model 5 (as Simplified Model 1, but no explicit dynamics of effector cells *E*^*’*^ ≡ 0)

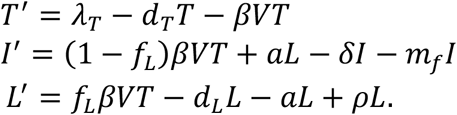

### S2 Model fits

Table S1 shows the comparisons of models under different assumptions. Figures S1-S6 (and Figure 1, main text) are the best fit of each model, with the population estimates provided in Tables S2-S6. Figures S7-S10 show the stratifications of best fit parameters for the Conway & Perelson model, Simplified Models 1, 2 and 3. Tables S7-13 give the individual estimates for the best fit of the Conway & Perelson model and Simplified Models 1 (with and without covariate on *K*_*B*_) through 5.

**Table S1.** Model fit comparison (Monolix 2023R1, Lixoft, SA, Antony, France). There are four categories separated by double-lines. Bolded values are the lowest values in the category. The first category contains the Conway and Perelson model with five variations without covariates. The second category tests single covariates for the best fit model in the first category. The third category shows two examples of using two covariates that produce the lowest fitting error. The last category tests the Simplified Model 1 without random effect for K_*B*_. Without a random effect, the population estimate for *K*_*B*_ is 16.88 (S.E. 0.12) cells mL^−1^.

| Model | Effector cell exhaustion | Dynamic latent reservoir | Dynamic effector cells | -2LL | BICc | Figure (Best-fit) |
| --- | --- | --- | --- | --- | --- | --- |
| Conway & Perelson | Yes | Yes | Yes | 547.38 | 625.29 | S1 |
| Simplified Model 1 | No | Yes | Yes | **545.33** | **605.22** | 1 (main) |
| Simplified Model 2 | No | No | Yes | 548.12 | 608.02 | S3 |
| Simplified Model 3 | Yes | No | Yes | 545.66 | 623.57 | S4 |
| Simplified Model 4 |  | No | No | 894.15 | 927.01 | S5 |
| Simplified Model 5 |  | Yes | No | 893.17 | 926.04 | S6 |
| Simplified Model 1 (covariate on $K_B$) | | | | **515.74** | **578.81** | S2 |
| Simplified Model 1 (covariate on $\beta$) | | | | 547.94 | 611.01 | — |
| Simplified Model 1 (covariate on $\lambda_E$) | | | | 546.29 | 609.36 | — |
| Simplified Model 1 (covariate on $b$) | | | | 532.59 | 595.66 | — |
| Simplified Model 1 (covariate on $p$) | | | | 546.98 | 610.05 | — |
| Simplified Model 1 (covariate on $m$) | | | | 543.33 | 606.40 | — |
| Simplified Model 1 (covariates on $K_B$ and $m$) | | | | **513.58** | **579.83** | — |
| Simplified Model 1 (covariates on $K_B$ and $b$) | | | | 514.84 | 581.09 | — |
| Simplified Model 1 (without random effect for $K_B$) | | | | 593.83 | 650.55 | — |

**Figure S1.**
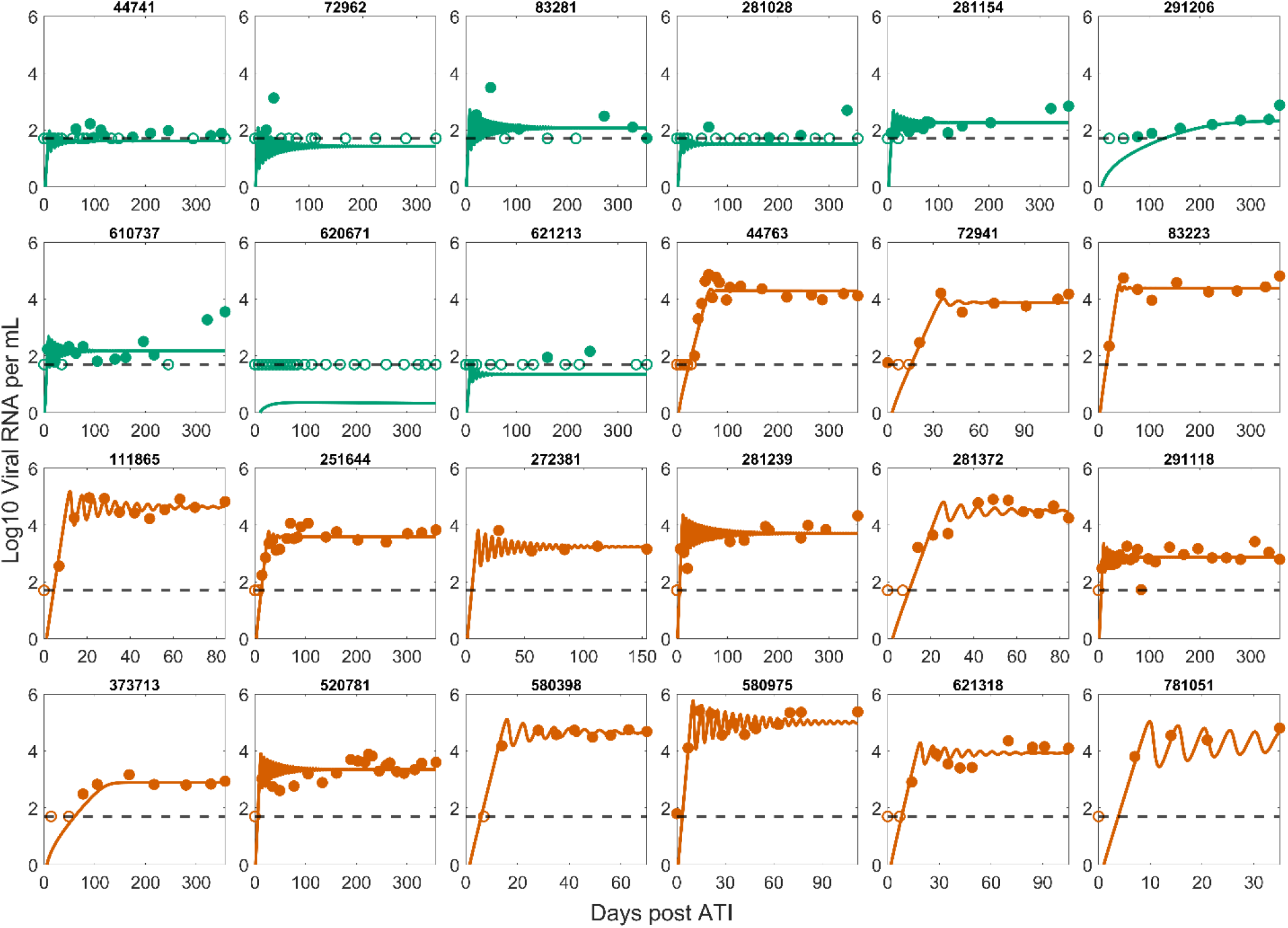
Best fit of the Conway & Perelson model to the post-ATI data from Sharaf et al. [57]. Green indicates PTC. Dark orange indicates NC. The horizontal dashed line is the limit of detection. Open circles are data points below the limit of detection (50 viral RNA copies/mL). Filled circles are data points above the limit of detection.

**Figure S2.**
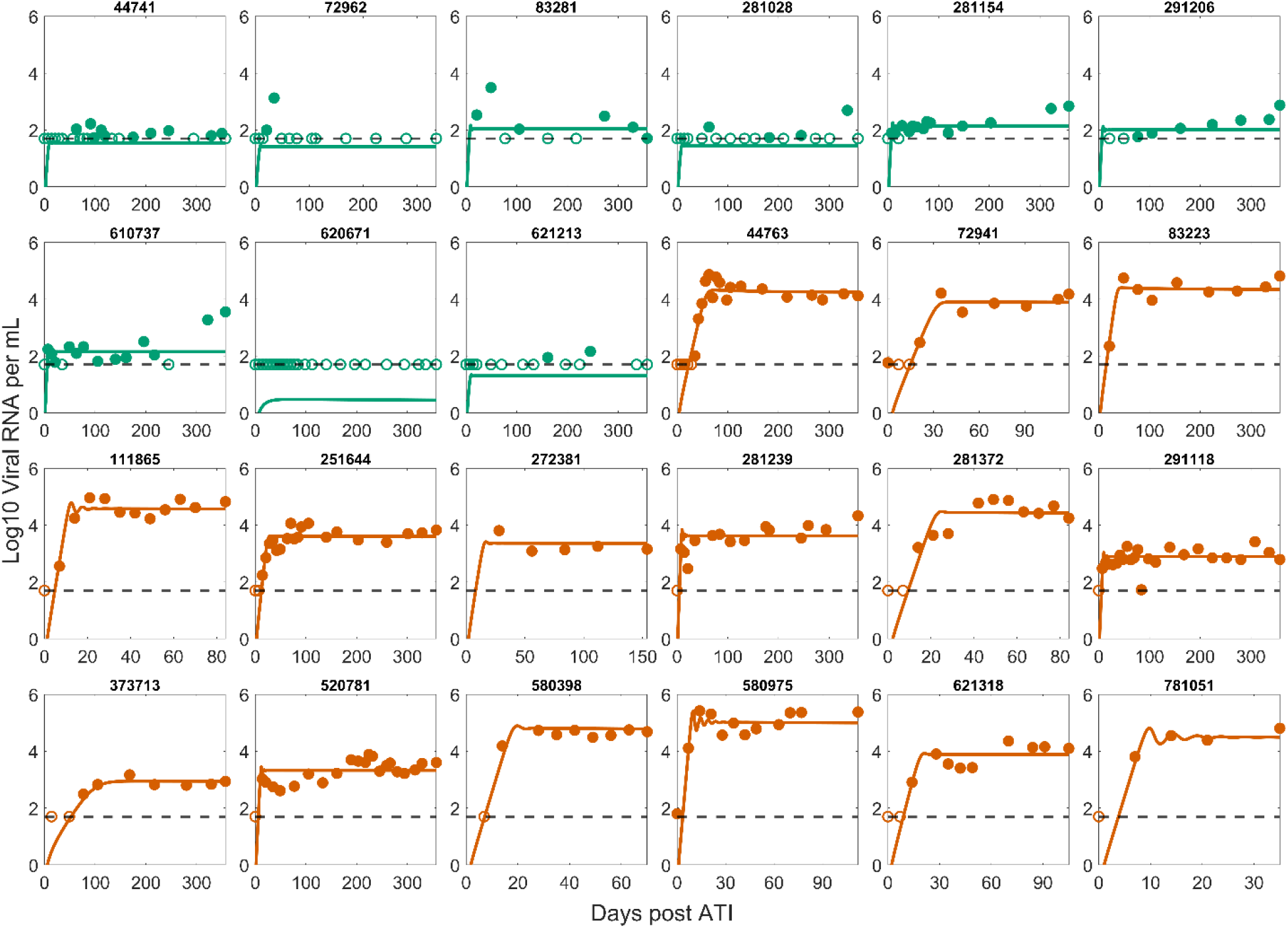
Best fit of Simplified Model 1 with a covariate on K_B_ to the post-ATI data from Sharaf et al. [57]. Lines show the model predictions using the best fit parameter values from Simplified Model 1 with a covariate on K_B_. Green indicates PTC. Dark orange indicates NC. The horizontal dashed line is the limit of detection (50 viral RNA copies/mL). Open circles are data points below the limit of detection. Filled circles are data points above the limit of detection.

**Figure S3.**
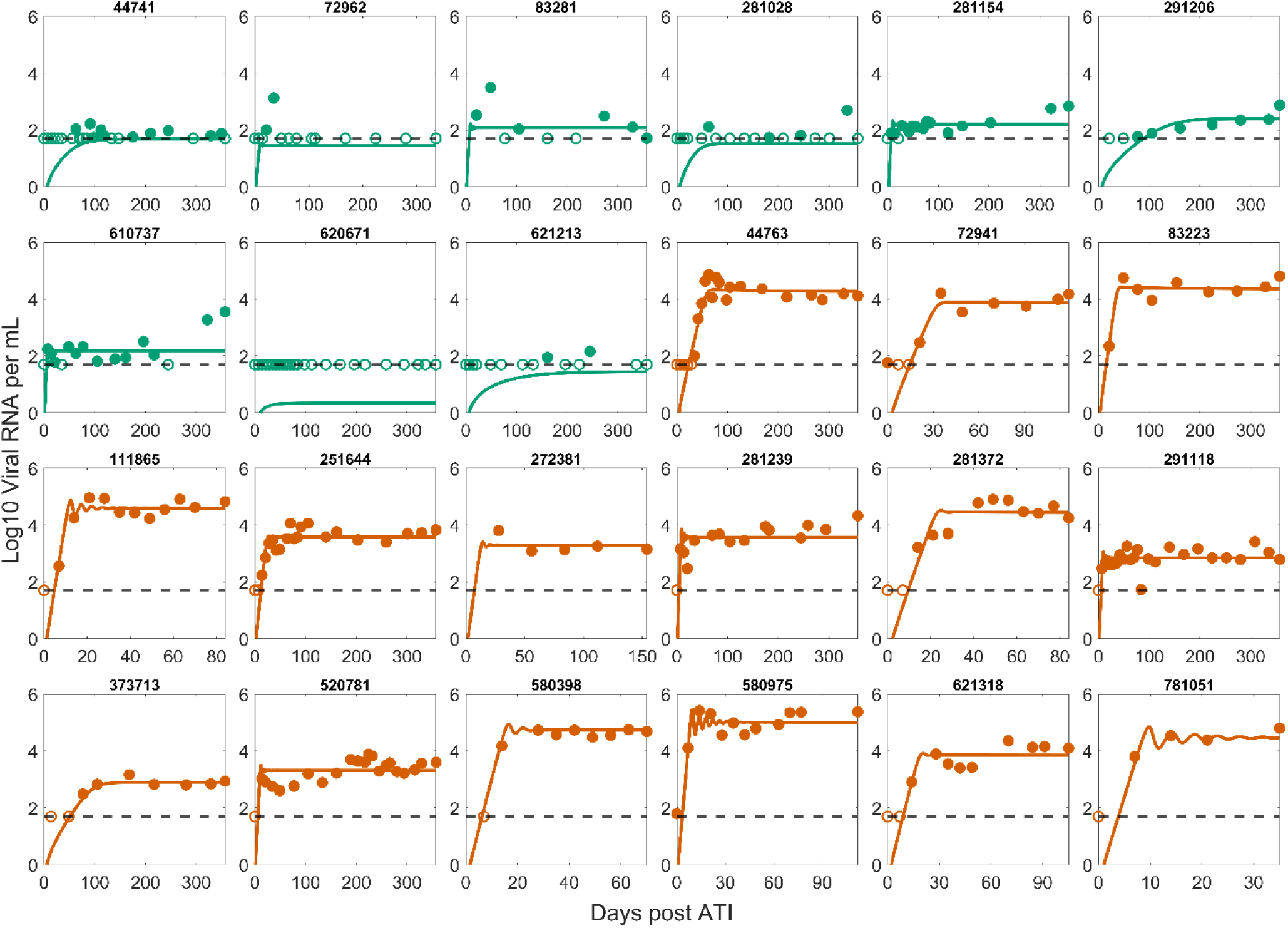
Best fit of the Simplified Model 2 to the post-ATI data from Sharaf et al. [57]. Green indicates PTC. Dark orange indicates NC. The horizontal dashed line is the limit of detection. Open circles are data points below the limit of detection (50 viral RNA copies/mL). Filled circles are data points above the limit of detection.

**Figure S4.**
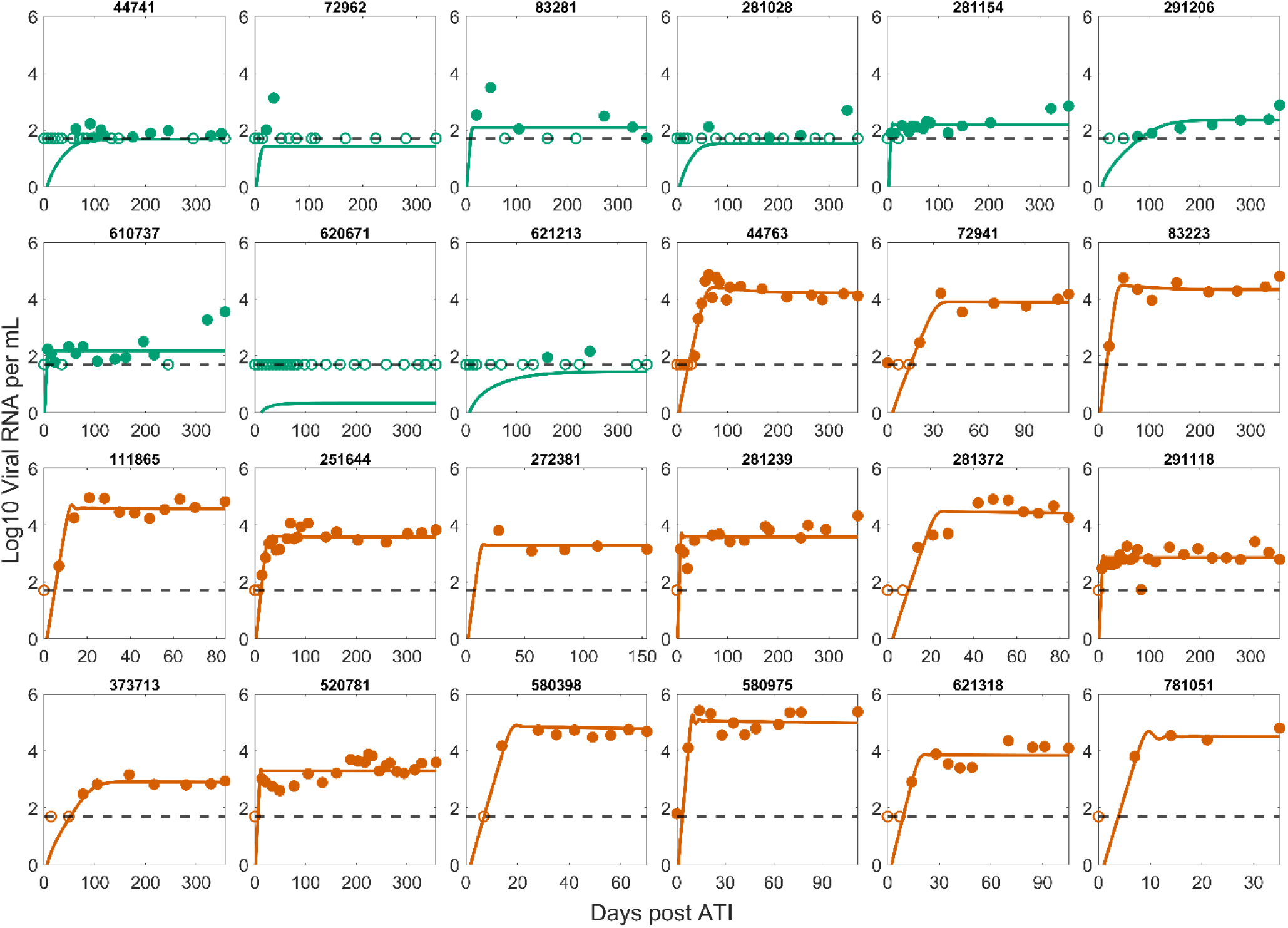
Best fit of the Simplified Model 3 to the post-ATI data from Sharaf et al. [57]. Green indicates PTC. Dark orange indicates NC. The horizontal dashed line is the limit of detection. Open circles are data points below the limit of detection (50 viral RNA copies/mL). Filled circles are data points above the limit of detection.

**Figure S5.**
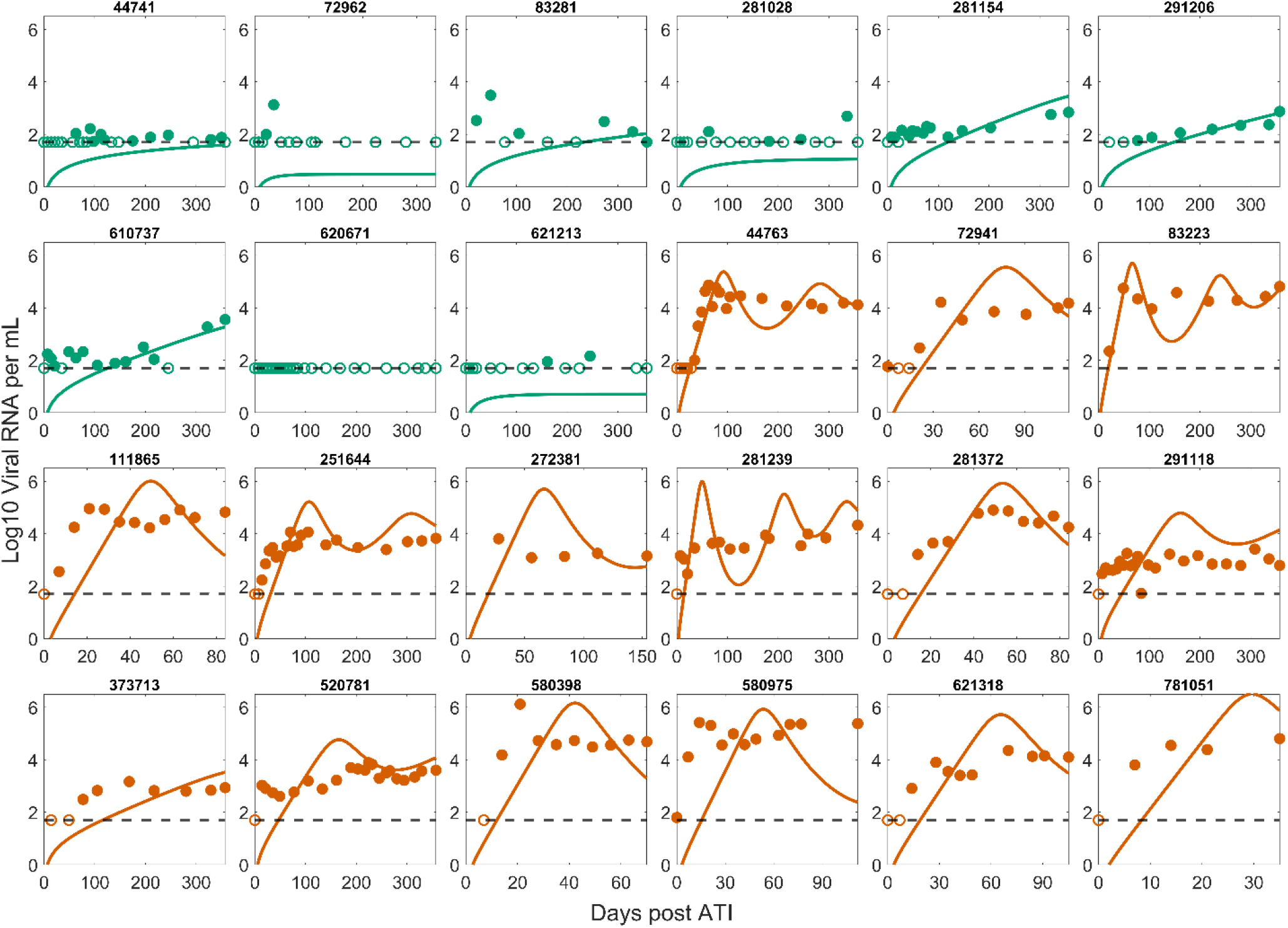
Best fit of the Simplified Model 4 to the post-ATI data from Sharaf et al. [57]. Green indicates PTC. Dark orange indicates NC. The horizontal dashed line is the limit of detection. Open circles are data points below the limit of detection (50 viral RNA copies/mL). Filled circles are data points above the limit of detection.

**Figure S6.**
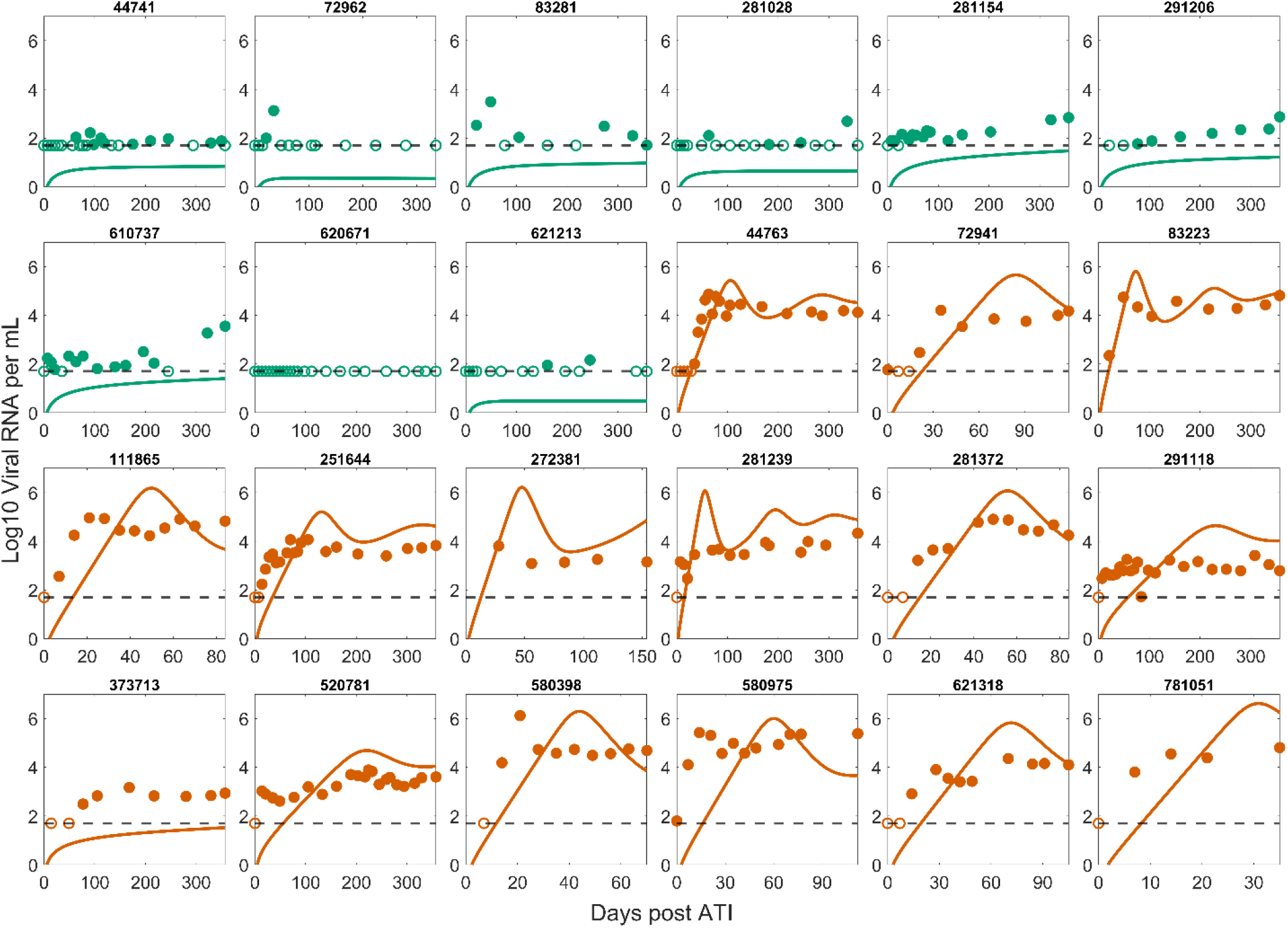
Best fit of the Simplified Model 5 to the post-ATI data from Sharaf et al. [57]. Green indicates PTC. Dark orange indicates NC. The horizontal dashed line is the limit of detection. Open circles are data points below the limit of detection (50 viral RNA copies/mL). Filled circles are data points above the limit of detection.

### S3 Stratification of best fit parameters

**Figure S7.**
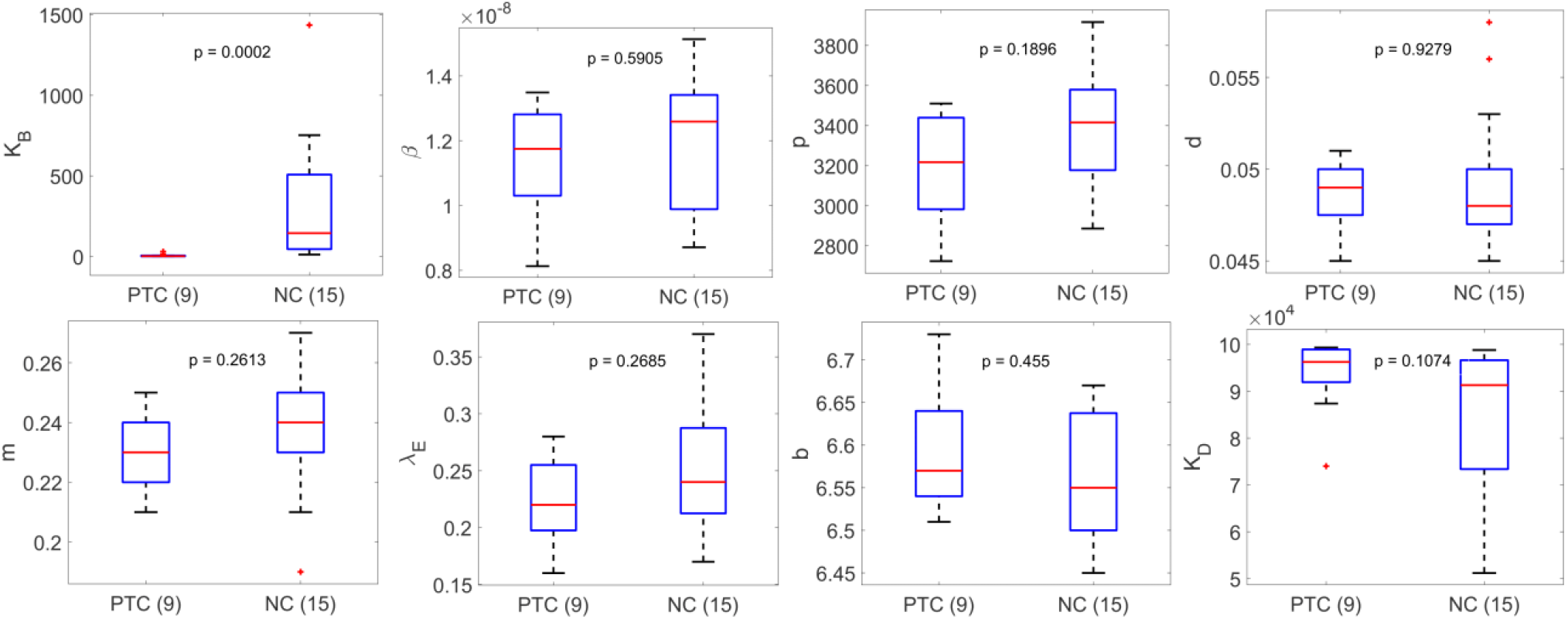
Summary of best-fit parameters in the Conway & Perelson Model stratified based on PTC or NC. The difference in the values of *K*_*B*_ is the most significant between PTC and NC.

**Figure S8.**
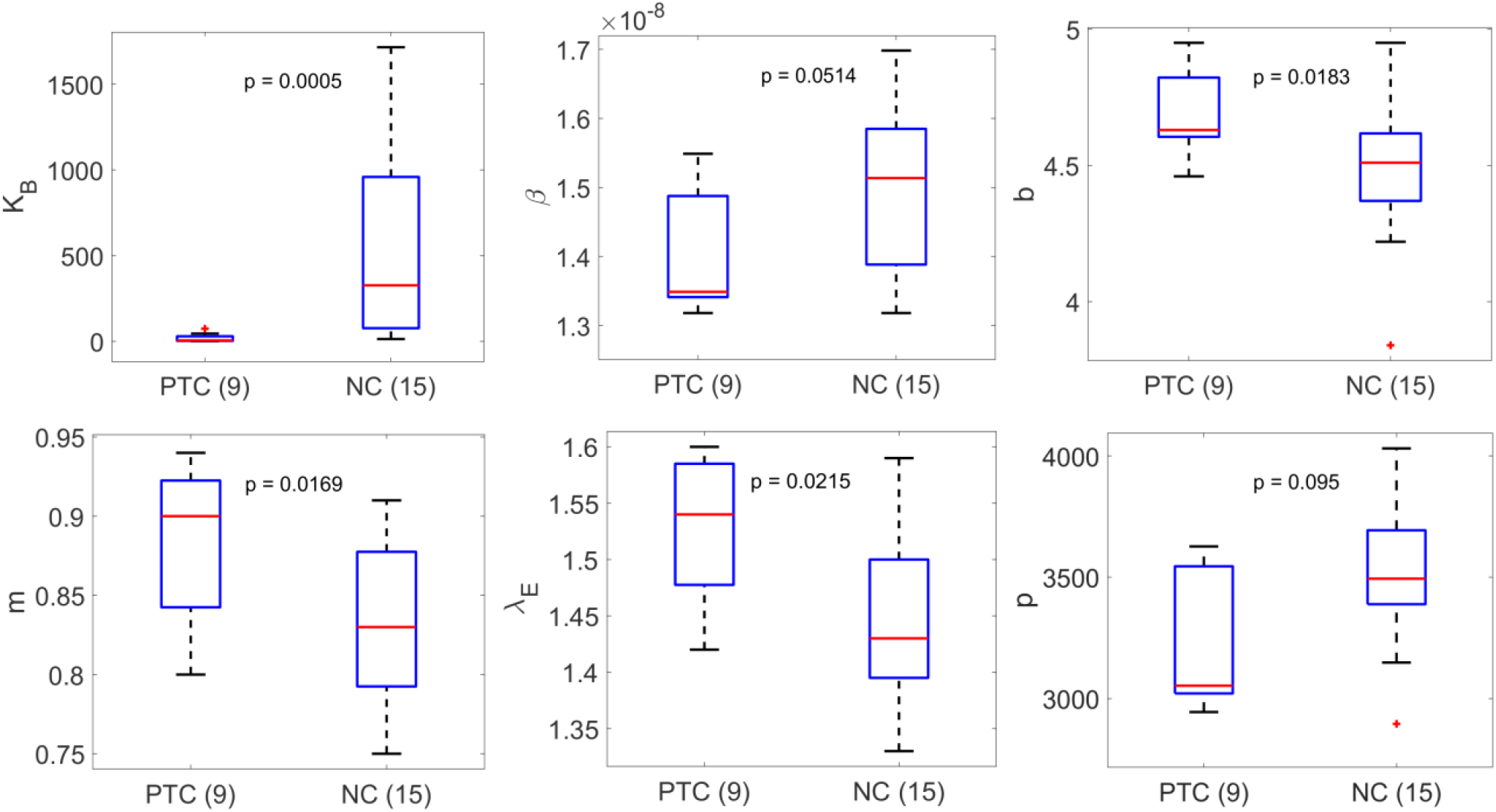
Summary of best fit parameters in Simplified Model 1 stratified based on PTC or NC. The difference in the values of K_B_ is the most significant between PTC and NC.

**Figure S9.**
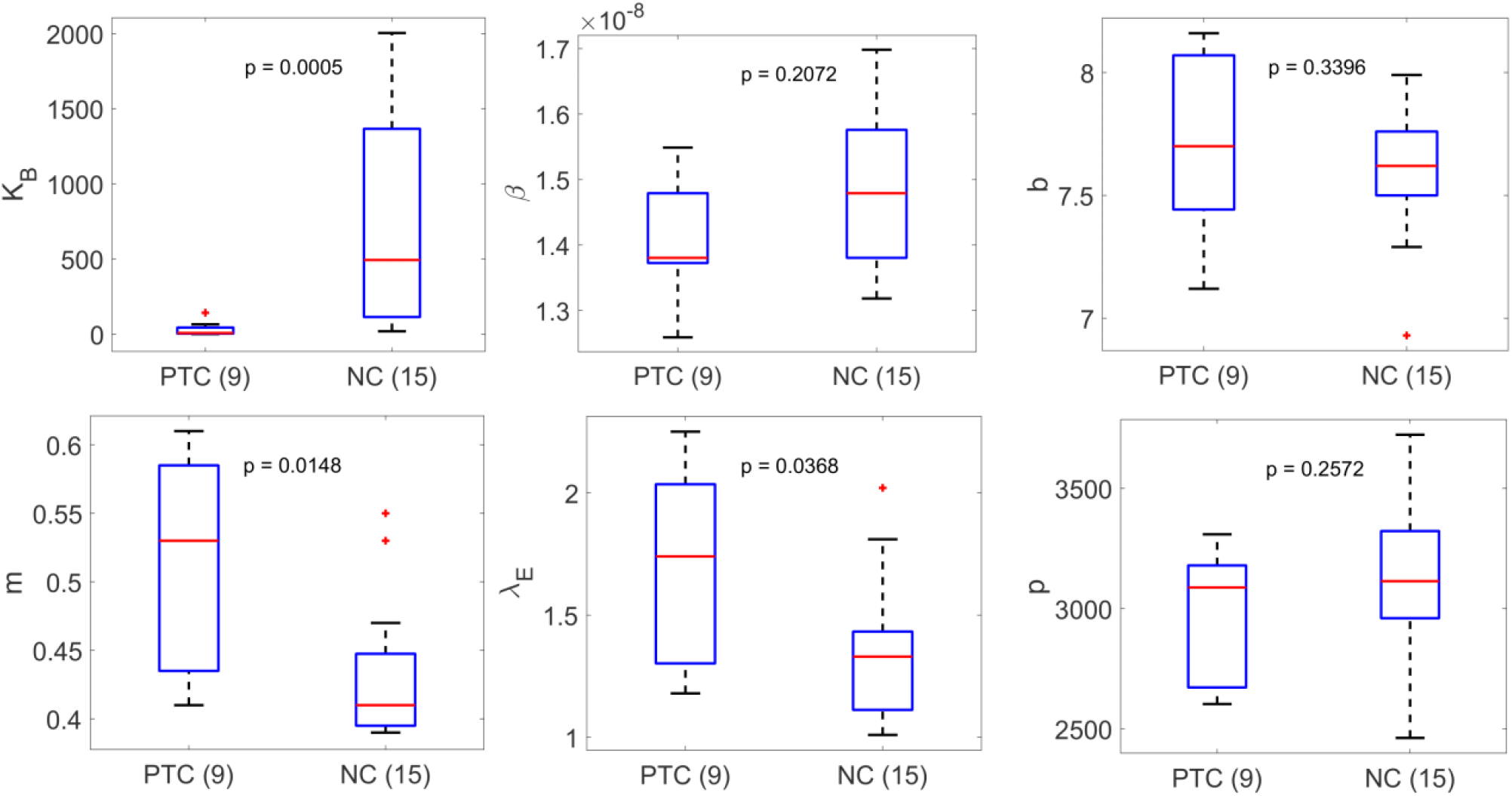
Summary of best-fit parameters in the Simplified Model 2 stratified based on PTC or NC. The difference in the values of *K*_*B*_ is the most significant between PTC and NC.

**Figure S10.**
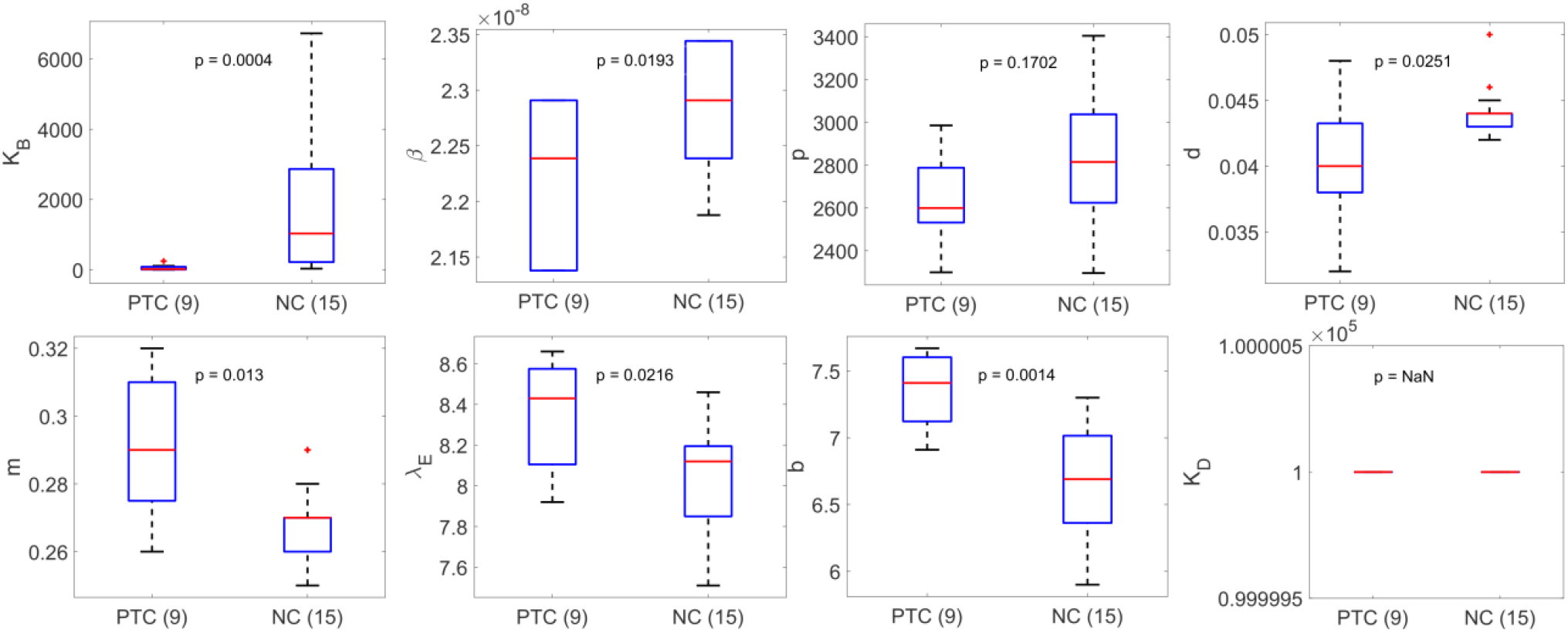
Summary of best-fit parameters in the Simplified Model 3 stratified based on PTC or NC. The difference in the values of *K*_*B*_ is the most significant between PTC and NC.

### S4 Best fit parameters for each individual

**Table S2.** Best-fit population parameters for the Conway & Perelson Model vs. reference values from Conway and Perelson [49]. Note the estimated values of *K*_*D*_ and *d* result in a negligible effect of exhaustion.

| *Parameter* | *Best fit values (population estimate)* | *Reference values* |
| --- | --- | --- |
| $\beta$ | $1.20 \times 10^{-8}$<br>$\text{mL (HIV RNA copies)}^{-1} \text{day}^{-1}$ | $1.5 \times 10^{-8}$<br>$\text{mL (HIV RNA copies)}^{-1} \text{day}^{-1}$ |
| $\lambda_E$ | $0.25 \text{ cells mL}^{-1} \text{day}^{-1}$ | $1 \text{ cells mL}^{-1} \text{day}^{-1}$ |
| $m$ | $0.24 \text{ mL cells}^{-1} \text{day}^{-1}$ | $0.42 \text{ mL cells}^{-1} \text{day}^{-1}$ |
| $b$ | $6.57 \text{ day}^{-1}$ | $1 \text{ day}^{-1}$ |
| $K_B$ | $29.35 \text{ cells mL}^{-1}$ | $0.1 \text{ cells mL}^{-1}$ |
| $p$ | $3337 \text{ (HIV RNA copies) day}^{-1}$ | $2000 \text{ (HIV RNA copies) day}^{-1}$ |
| $K_D$ | $95188 \text{ cells mL}^{-1}$ | $5 \text{ cells mL}^{-1}$ |
| $d$ | $0.05 \text{ day}^{-1}$ | $2 \text{ day}^{-1}$ |

**Table S3.** Best-fit population parameters for the Simplified Model 2 vs. reference values from Conway and Perelson [49].

| *Parameter* | *Best fit values (population estimate)* | *Reference values* |
| --- | --- | --- |
| $\beta$ | $1.51 \times 10^{-8}$<br>$\text{mL (HIV RNA copies)}^{-1} \text{day}^{-1}$ | $1.5 \times 10^{-8}$<br>$\text{mL (HIV RNA copies)}^{-1} \text{day}^{-1}$ |
| $\lambda_E$ | $1.33 \text{ cells mL}^{-1} \text{day}^{-1}$ | $1 \text{ cells mL}^{-1} \text{day}^{-1}$ |
| $m$ | $0.44 \text{ mL cells}^{-1} \text{day}^{-1}$ | $0.42 \text{ mL cells}^{-1} \text{day}^{-1}$ |
| $b$ | $7.66 \text{ day}^{-1}$ | $1 \text{ day}^{-1}$ |
| $K_B$ | $65.99 \text{ cells mL}^{-1}$ | $0.1 \text{ cells mL}^{-1}$ |
| $p$ | $3104 \text{ (HIV RNA copies) day}^{-1}$ | $2000 \text{ (HIV RNA copies) day}^{-1}$ |

**Table S4.** Best-fit population parameters for the Simplified Model 3 vs. reference values from Conway and Perelson [49]. Note that the estimated values of *K*_*D*_ and *d* together imply the exhaustion effect is very small, which may be because of this particular set of participants or that it cannot be observed from the limited data.

| *Parameter* | *Best fit values (population estimate)* | *Reference values* |
| --- | --- | --- |
| $\beta$ | $2.29 \times 10^{-8}$<br>$\text{mL (HIV RNA copies)}^{-1} \text{day}^{-1}$ | $1.5 \times 10^{-8}$<br>$\text{mL (HIV RNA copies)}^{-1} \text{day}^{-1}$ |
| $\lambda_E$ | $8.06 \text{ cells mL}^{-1} \text{day}^{-1}$ | $1 \text{ cells mL}^{-1} \text{day}^{-1}$ |
| $m$ | $0.27 \text{ mL cells}^{-1} \text{day}^{-1}$ | $0.42 \text{ mL cells}^{-1} \text{day}^{-1}$ |
| $b$ | $6.8 \text{ day}^{-1}$ | $1 \text{ day}^{-1}$ |
| $K_B$ | $122.97 \text{ cells mL}^{-1}$ | $0.1 \text{ cells mL}^{-1}$ |
| $p$ | $2798 \text{ (HIV RNA copies) day}^{-1}$ | $2000 \text{ (HIV RNA copies) day}^{-1}$ |
| $K_D$ | $100000 \text{ cells mL}^{-1}$ | $5 \text{ cells mL}^{-1}$ |
| $d$ | $0.044 \text{ day}^{-1}$ | $2 \text{ day}^{-1}$ |

**Table S5.** Best-fit population parameters for the Simplified Model 4 vs. reference values from Conway and Perelson [49]. Note that *m*^*^ is not the same as *m*.

| *Parameter* | *Best fit values (population estimate)* | *Reference values* |
| --- | --- | --- |
| $\beta$ | $2.04 \times 10^{-8}$<br>$mL (HIV RNA copies)^{-1} day^{-1}$ | $1.5 \times 10^{-8}$<br>$mL (HIV RNA copies)^{-1} day^{-1}$ |
| $m^*$ | $1.4 mL day^{-1}$ | |
| $p$ | $2855 (HIV RNA copies) day^{-1}$ | $2000 (HIV RNA copies) day^{-1}$ |

**Table S6.** Best-fit population parameters for the Simplified Model 5 vs. reference values from Conway and Perelson [49]. Note that *m*^*^ is not the same as *m*.

| *Parameter* | *Best fit values (population estimate)* | *Reference values* |
| --- | --- | --- |
| $\beta$ | $1.48 \times 10^{-8}$<br>$mL (HIV RNA copies)^{-1} day^{-1}$ | $1.5 \times 10^{-8}$<br>$mL (HIV RNA copies)^{-1} day^{-1}$ |
| $m^*$ | $1.18 mL day^{-1}$ | |
| $p$ | $3608 (HIV RNA copies) day^{-1}$ | $2000 (HIV RNA copies) day^{-1}$ |

**Table S7.**
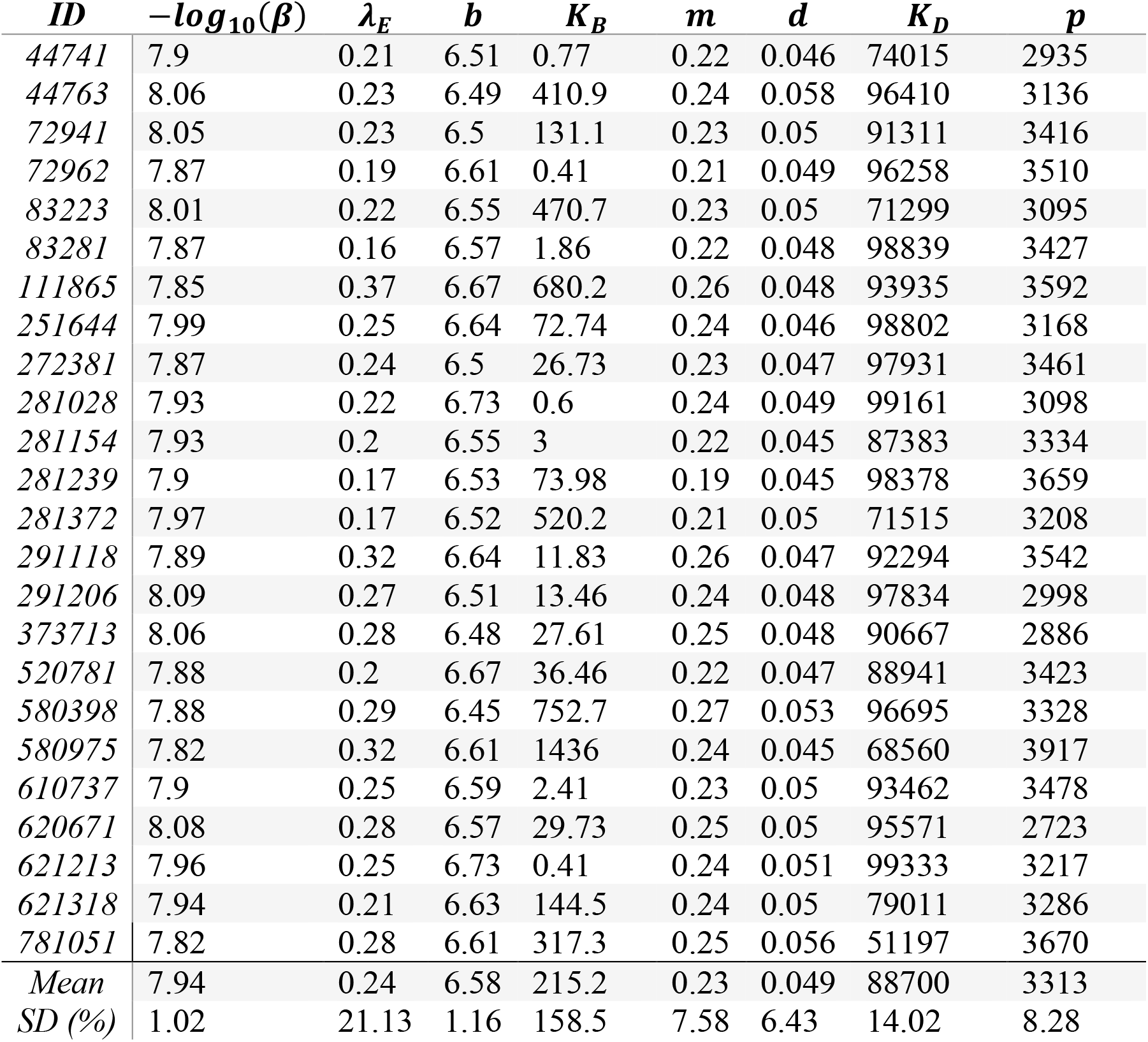
Individual best-fit parameters – Conway & Perelson model.

**Table S8.** Individual best-fit parameters – Simplified Model 1.

| *ID* | $-\log_{10}(\beta)$ | $\lambda_E$ | $b$ | $K_B$ | $m$ | $p$ |
| --- | --- | --- | --- | --- | --- | --- |
| 44741 | 7.87 | 1.6 | 4.63 | 14.67 | 0.92 | 3053 |
| 44763 | 7.85 | 1.43 | 4.67 | 1716 | 0.87 | 2896 |
| 72941 | 7.87 | 1.55 | 4.22 | 326.9 | 0.89 | 3422 |
| 72962 | 7.85 | 1.49 | 4.95 | 0.99 | 0.9 | 3468 |
| 83223 | 7.86 | 1.51 | 4.23 | 1222 | 0.88 | 3225 |
| 83281 | 7.83 | 1.49 | 4.78 | 2.95 | 0.85 | 3536 |
| 111865 | 7.8 | 1.38 | 4.43 | 568.4 | 0.78 | 3707 |
| 251644 | 7.86 | 1.54 | 4.63 | 149 | 0.89 | 3463 |
| 272381 | 7.82 | 1.46 | 4.58 | 42.88 | 0.83 | 3494 |
| 281028 | 7.87 | 1.58 | 4.61 | 5.08 | 0.91 | 3053 |
| 281154 | 7.82 | 1.44 | 4.95 | 3.33 | 0.8 | 3570 |
| 281239 | 7.79 | 1.39 | 4.35 | 52.61 | 0.79 | 3787 |
| 281372 | 7.84 | 1.47 | 4.43 | 795.3 | 0.84 | 3379 |
| 291118 | 7.8 | 1.41 | 4.95 | 13.41 | 0.8 | 3651 |
| 291206 | 7.88 | 1.56 | 4.46 | 73.62 | 0.89 | 3040 |
| 373713 | 7.88 | 1.59 | 4.64 | 161.2 | 0.91 | 3148 |
| 520781 | 7.81 | 1.41 | 4.51 | 34.97 | 0.81 | 3650 |
| 580398 | 7.82 | 1.43 | 4.52 | 1422 | 0.81 | 3494 |
| 580975 | 7.77 | 1.33 | 3.84 | 1014 | 0.75 | 4031 |
| 610737 | 7.81 | 1.42 | 4.68 | 2.92 | 0.82 | 3627 |
| 620671 | 7.88 | 1.6 | 4.59 | 45.02 | 0.94 | 2964 |
| 621213 | 7.87 | 1.54 | 4.62 | 24.08 | 0.93 | 2945 |
| 621318 | 7.84 | 1.47 | 4.51 | 191.5 | 0.84 | 3418 |
| 781051 | 7.77 | 1.34 | 4.52 | 411.5 | 0.75 | 3881 |
| *Mean* | 7.84 | 1.48 | 4.55 | 345.6 | 0.85 | 3413 |
| *SD (%)* | 0.43 | 5.37 | 5.29 | 144.1 | 6.51 | 8.95 |

**Table S9.** Individual best-fit parameters – Simplified Model 1 with a covariate on *K*_*B*_ (1 is PTC and 2 is NC). For *K*_*B*_, the mean and SD (%) are reported for individual group PTC/NC.

| *ID* | $-\log_{10}(\beta)$ | $\lambda_E$ | $b$ | $K_B$ | $m$ | $p$ | *covariate* |
| --- | --- | --- | --- | --- | --- | --- | --- |
| 44741 | 7.82 | 0.54 | 5.23 | 0.94 | 1.69 | 3097 | 1 |
| 44763 | 7.85 | 0.52 | 5.13 | 1488 | 1.72 | 2657 | 2 |
| 72941 | 7.87 | 0.6 | 5.15 | 359.8 | 1.76 | 3123 | 2 |
| 72962 | 7.83 | 0.54 | 5.75 | 0.78 | 1.69 | 3179 | 1 |
| 83223 | 7.86 | 0.55 | 4.25 | 878.9 | 1.72 | 2977 | 2 |
| 83281 | 7.8 | 0.48 | 5.09 | 2.17 | 1.69 | 3261 | 1 |
| 111865 | 7.78 | 0.45 | 4.97 | 603.1 | 1.65 | 3424 | 2 |
| 251644 | 7.85 | 0.58 | 5.35 | 160.7 | 1.74 | 3062 | 2 |
| 272381 | 7.81 | 0.54 | 6.18 | 75.81 | 1.71 | 3104 | 2 |
| 281028 | 7.82 | 0.54 | 5.14 | 0.74 | 1.66 | 3112 | 1 |
| 281154 | 7.8 | 0.48 | 5.29 | 2.89 | 1.7 | 3253 | 1 |
| 281239 | 7.77 | 0.47 | 5.78 | 80.96 | 1.66 | 3466 | 2 |
| 281372 | 7.83 | 0.52 | 4.61 | 699.7 | 1.73 | 3066 | 2 |
| 291118 | 7.78 | 0.51 | 6.41 | 20.21 | 1.75 | 3331 | 2 |
| 291206 | 7.8 | 0.49 | 4.85 | 1.86 | 1.69 | 3276 | 1 |
| 373713 | 7.86 | 0.63 | 5.26 | 201.7 | 1.92 | 2764 | 2 |
| 520781 | 7.79 | 0.51 | 6.24 | 55.49 | 1.7 | 3241 | 2 |
| 580398 | 7.82 | 0.48 | 4.2 | 1104 | 1.7 | 3179 | 2 |
| 580975 | 7.76 | 0.42 | 4.45 | 1125 | 1.6 | 3749 | 2 |
| 610737 | 7.79 | 0.47 | 5.38 | 2.74 | 1.68 | 3351 | 1 |
| 620671 | 7.86 | 0.66 | 5.49 | 0.92 | 1.86 | 2722 | 1 |
| 621213 | 7.82 | 0.55 | 5.67 | 0.64 | 1.71 | 3078 | 1 |
| 621318 | 7.83 | 0.55 | 5.41 | 227.7 | 1.67 | 3106 | 2 |
| 781051 | 7.76 | 0.43 | 5.24 | 455.6 | 1.63 | 3613 | 2 |
| *Mean* | 7.82 | 0.52 | 5.27 | 1.52/502.4 | 1.71 | 3175 |  |
| *SD (%)* | 0.41 | 11.07 | 10.57 | 56.04/89.14 | 3.84 | 7.85 |  |

**Table S10.** Individual best-fit parameters – Simplified Model 2.

| *ID* | $-\log_{10}(\beta)$ | $\lambda_E$ | *b* | $K_B$ | *m* | *p* |
| --- | --- | --- | --- | --- | --- | --- |
| 44741 | 7.86 | 1.89 | 8.06 | 21.76 | 0.6 | 2681 |
| 44763 | 7.86 | 1.41 | 7.99 | 1950 | 0.41 | 2462 |
| 72941 | 7.88 | 1.81 | 7.48 | 493.7 | 0.53 | 3127 |
| 72962 | 7.85 | 1.55 | 8.16 | 1.24 | 0.47 | 3088 |
| 83223 | 7.86 | 1.52 | 6.93 | 1439 | 0.47 | 2788 |
| 83281 | 7.83 | 1.31 | 7.94 | 3.81 | 0.42 | 3177 |
| 111865 | 7.8 | 1.1 | 7.62 | 971.2 | 0.39 | 3341 |
| 251644 | 7.88 | 1.33 | 7.78 | 179.1 | 0.44 | 2955 |
| 272381 | 7.83 | 1.34 | 7.66 | 64.83 | 0.44 | 3085 |
| 281028 | 7.86 | 1.74 | 7.15 | 7.44 | 0.54 | 2603 |
| 281154 | 7.83 | 1.28 | 8.1 | 5.15 | 0.44 | 3187 |
| 281239 | 7.79 | 1.15 | 7.81 | 92.8 | 0.39 | 3406 |
| 281372 | 7.85 | 1.44 | 7.56 | 1147 | 0.44 | 2975 |
| 291118 | 7.81 | 1.21 | 7.81 | 19.49 | 0.41 | 3267 |
| 291206 | 7.86 | 1.99 | 7.54 | 142.6 | 0.53 | 2648 |
| 373713 | 7.87 | 2.02 | 7.58 | 235.1 | 0.55 | 2762 |
| 520781 | 7.82 | 1.2 | 7.7 | 58.35 | 0.41 | 3220 |
| 580398 | 7.82 | 1.01 | 7.58 | 1586 | 0.41 | 3114 |
| 580975 | 7.77 | 1.04 | 7.29 | 2004 | 0.39 | 3723 |
| 610737 | 7.81 | 1.18 | 7.7 | 4.1 | 0.41 | 3309 |
| 620671 | 7.87 | 2.17 | 7.66 | 65.31 | 0.58 | 2694 |
| 621213 | 7.9 | 2.25 | 7.12 | 37.28 | 0.61 | 3122 |
| 621318 | 7.85 | 1.41 | 7.47 | 266.1 | 0.45 | 3046 |
| 781051 | 7.77 | 1.08 | 7.67 | 655.7 | 0.39 | 3561 |
| *Mean* | 7.84 | 1.48 | 7.64 | 477.1 | 0.46 | 3056 |
| *SD (%)* | 0.43 | 24.59 | 3.94 | 135.8 | 15.01 | 10.03 |

**Table S11.**
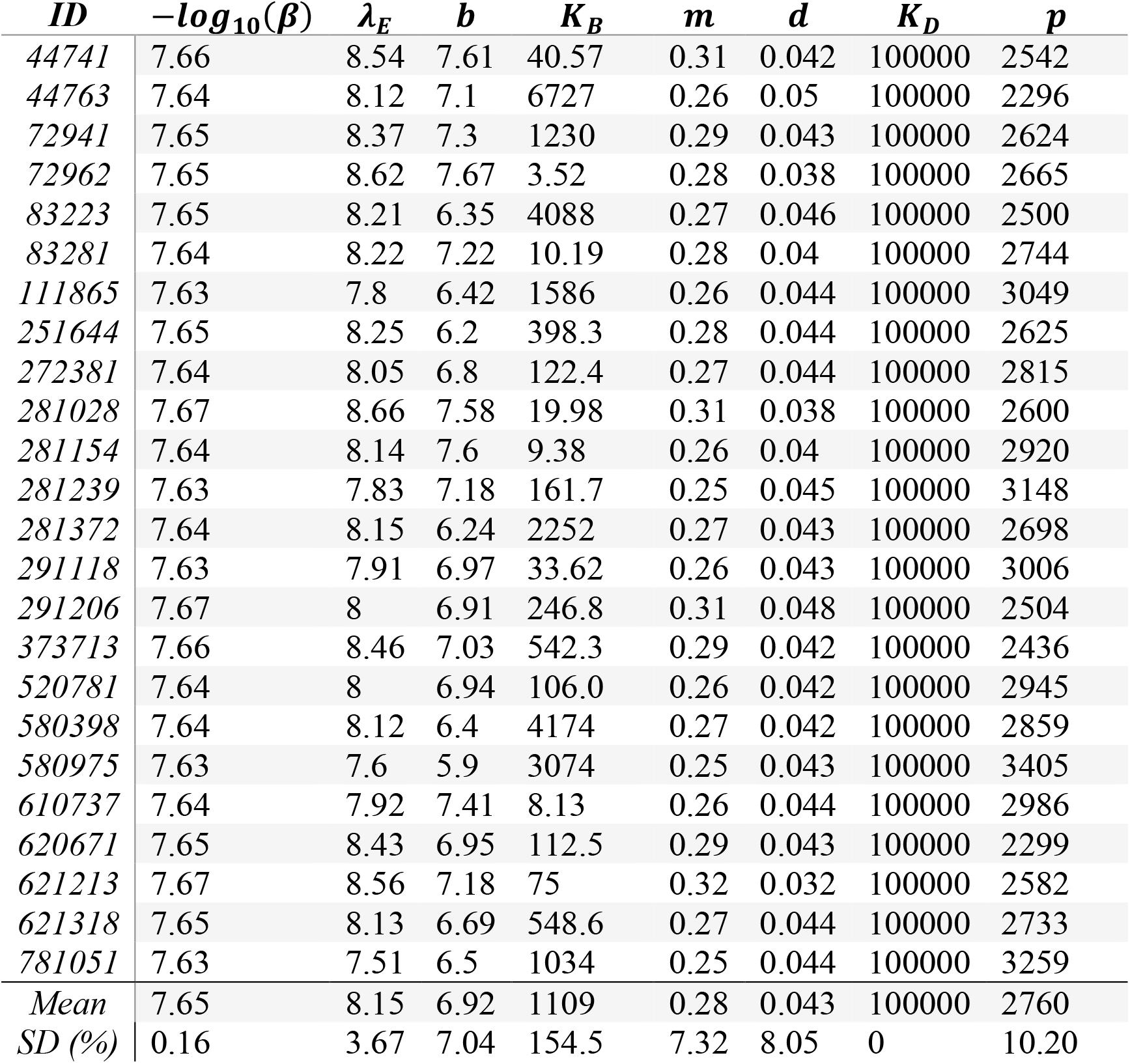
Individual best-fit parameters – Simplified Model 3.

**Table S12.** Individual best-fit parameters – Simplified Model 4.

| *ID* | $-\log_{10}(\beta)$ | *m* | *p* |
| --- | --- | --- | --- |
| 44741 | 7.69 | 1.44 | 2767 |
| 44763 | 7.69 | 1.38 | 2855 |
| 72941 | 7.68 | 1.38 | 2886 |
| 72962 | 7.69 | 1.46 | 2744 |
| 83223 | 7.68 | 1.36 | 2901 |
| 83281 | 7.69 | 1.45 | 2775 |
| 111865 | 7.68 | 1.33 | 2962 |
| 251644 | 7.68 | 1.41 | 2853 |
| 272381 | 7.68 | 1.37 | 2911 |
| 281028 | 7.69 | 1.45 | 2759 |
| 281154 | 7.69 | 1.45 | 2790 |
| 281239 | 7.68 | 1.32 | 2951 |
| 281372 | 7.68 | 1.35 | 2958 |
| 291118 | 7.69 | 1.44 | 2836 |
| 291206 | 7.69 | 1.44 | 2775 |
| 373713 | 7.69 | 1.44 | 2782 |
| 520781 | 7.69 | 1.44 | 2841 |
| 580398 | 7.68 | 1.32 | 3020 |
| 580975 | 7.68 | 1.34 | 2958 |
| 610737 | 7.69 | 1.44 | 2787 |
| 620671 | 7.69 | 1.5 | 2669 |
| 621213 | 7.69 | 1.45 | 2751 |
| 621318 | 7.68 | 1.37 | 2914 |
| 781051 | 7.68 | 1.27 | 3148 |
| *Mean* | 7.69 | 1.4 | 2858 |
| *SD (%)* | 0.065 | 4.08 | 3.69 |

**Table S13.** Individual best-fit parameters – Simplified Model 5.

| *ID* | $-\log_{10}(\beta)$ | *m* | *p* |
| --- | --- | --- | --- |
| 44741 | 7.84 | 1.21 | 3519 |
| 44763 | 7.83 | 1.17 | 3607 |
| 72941 | 7.83 | 1.17 | 3643 |
| 72962 | 7.84 | 1.22 | 3496 |
| 83223 | 7.83 | 1.14 | 3643 |
| 83281 | 7.84 | 1.21 | 3531 |
| 111865 | 7.82 | 1.13 | 3725 |
| 251644 | 7.83 | 1.2 | 3612 |
| 272381 | 7.81 | 1.14 | 3745 |
| 281028 | 7.84 | 1.21 | 3512 |
| 281154 | 7.84 | 1.21 | 3547 |
| 281239 | 7.82 | 1.14 | 3687 |
| 281372 | 7.82 | 1.14 | 3699 |
| 291118 | 7.83 | 1.21 | 3554 |
| 291206 | 7.84 | 1.21 | 3529 |
| 373713 | 7.84 | 1.21 | 3536 |
| 520781 | 7.83 | 1.21 | 3571 |
| 580398 | 7.81 | 1.13 | 3762 |
| 580975 | 7.82 | 1.15 | 3706 |
| 610737 | 7.84 | 1.21 | 3544 |
| 620671 | 7.85 | 1.24 | 3414 |
| 621213 | 7.84 | 1.22 | 3502 |
| 621318 | 7.82 | 1.16 | 3664 |
| 781051 | 7.8 | 1.1 | 3861 |
| *Mean* | 7.83 | 1.18 | 3609 |
| *SD (%)* | 0.15 | 3.21 | 2.87 |

### S5 Analytical approximation of the early viral rebound dynamics

In order to better explain why *K*_*B*_ separates the NC and PTC participants, we approximate analytically the viral set point after rebound for Simplified Model 1:

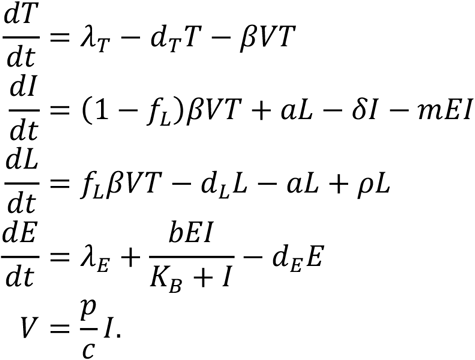

We are interested in the viral set point attained < 1 year post-ATI and as HIV-1 infection progresses slowly, we approximate the target cell population by its disease-free equilibrium *T* = *λ*_*T*_/*d*_*T*_. It is worth pointing out that PWH, who experienced rapid viral rebound and potentially rapid decline of target cells, re-started ART. This approximation also works for the period before ART.

In this deterministic model, the contribution from latent cells to the viral rebound dynamics (e.g., the rate that virus becomes detectable and the magnitude of the set point viral load) is negligible. Thus, we assume that *L* = 0 for this approximation, which will simplify the algebraic details further. Together, these considerations result in the following transient system which approximates the early viral rebound dynamics of the full model:

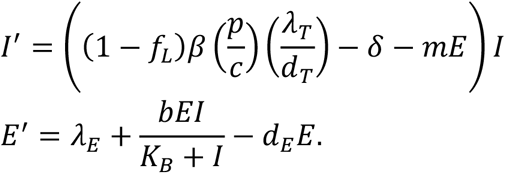

This approximated system with one infected cell compartment and one effector cell compartment is inspired by a previous study (Baral et al., 2019). Now, solving for *E*^′^ = 0 gives

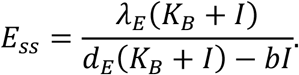

Substituting this expression for *E* into the *I*′ equation,

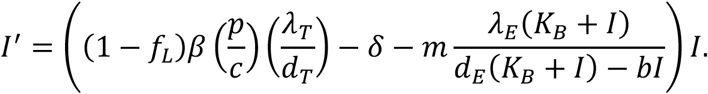

Setting *I*^′^ = 0 and solving for the non-zero steady state *I*_*ss*_ gives

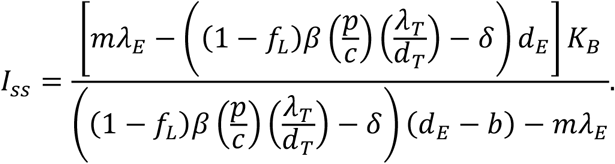

The corresponding critical value *V*_*c*_ for *V*(*t*) is given by

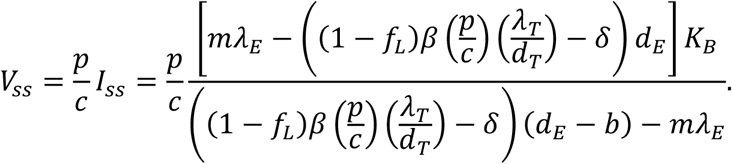

Figure S11 shows a comparison of *V*_*ss*_ with the predicted viral load set-point obtained with the full model dynamics.

**Figure S11.**
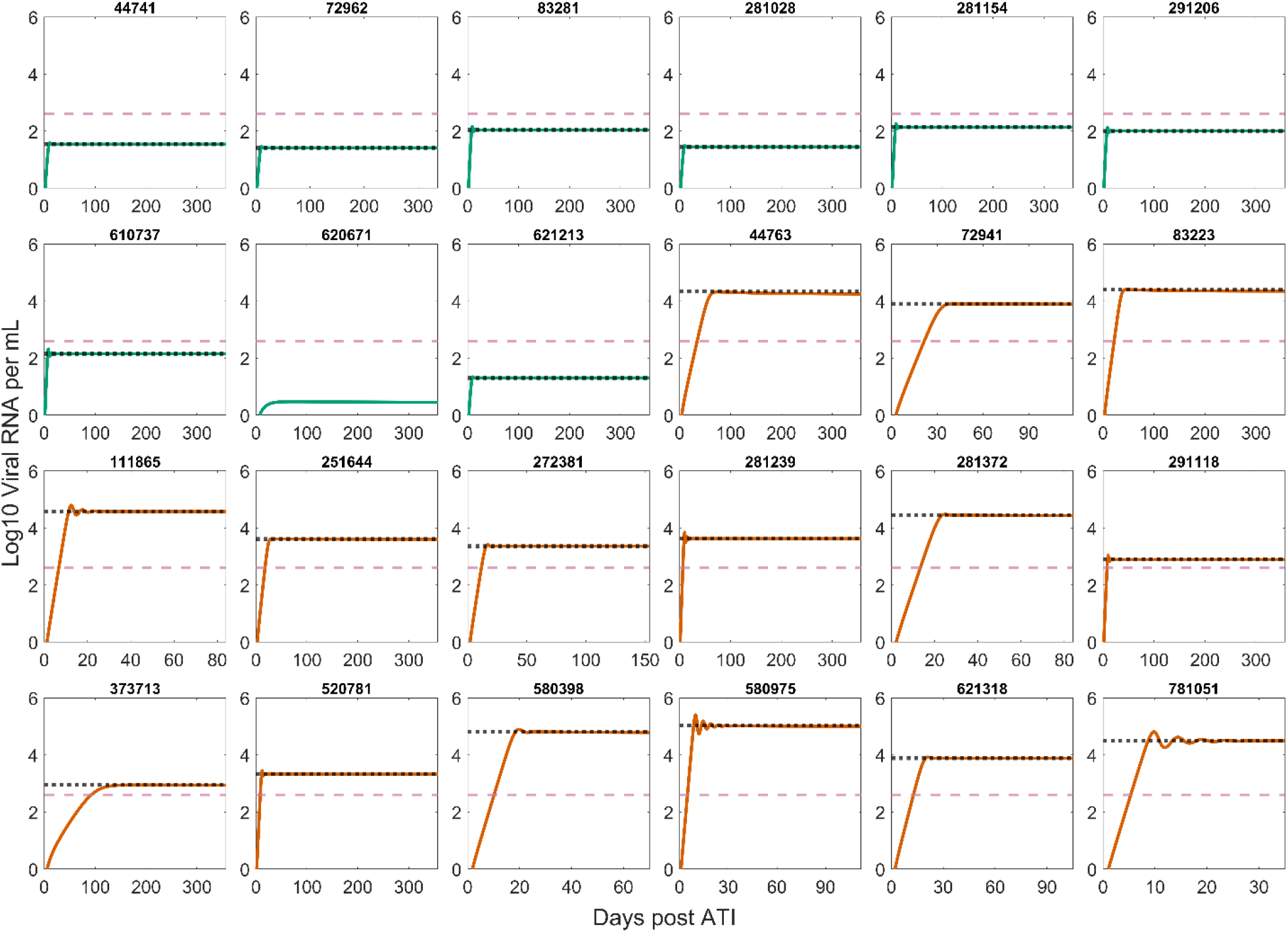
Approximation of the set point viral load V_ss_ using the best fit parameters for each participant compared with the numerical solution of Simplified Model 1 (Eqn. 2). The horizontal black dotted line is the approximation V_ss_ (Eqn. 3). Green and dark orange curves correspond to the viral load of the PTC and NC participants, respectively, as predicted by the Simplified Model 1. The horizontal pink dashed line is the 400 viral RNA copies/mL threshold used in the classification of PTC. For participant number 620671, V_ss_ is at 0.61 viral RNA copies/mL, which is the full model predicted viral set point after day ∼12000.

To arrive at this approximation of the viral set point, we made several crude simplifications, where the assumptions of limited exhaustion 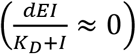 and constant target cells 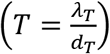 resulted in the removal of one of the two non-extinction steady states of the full model. However, it is not clear whether or when both steady states are positive since the algebraic calculation is fairly complicated for the full model. The model also contains a disease-free steady state, which is unstable assuming the baseline immune response is not sufficient to clear the virus on its own.

